# The development of ecological systems along paths of least resistance

**DOI:** 10.1101/2024.06.24.600194

**Authors:** Jie Deng, Otto X. Cordero, Tadashi Fukami, Simon A. Levin, Robert M. Pringle, Ricard Solé, Serguei Saavedra

## Abstract

A long-standing question in biology is whether there are common principles that characterize the development of ecological systems (the appearance of a group of taxa), regardless of organismal diversity and environmental context. Classic ecological theory holds that these systems develop following a sequenced orderly process that generally proceeds from fast-growing to slow-growing taxa and depends on life-history trade-offs. However, it is also possible that this developmental order is simply the path with the least environmental resistance for survival of the component species and hence favored by probability alone. Here, we use theory and data to show that the order from fast-to slow-growing taxa is the most likely developmental path for diverse systems when local taxon interactions self-organize to minimize environmental resistance. First, we demonstrate theoretically that a sequenced development is more likely than a simultaneous one, at least until the number of iterations becomes so large as to be ecologically implausible. We then show that greater diversity of taxa and life histories improves the likelihood of a sequenced order from fast-to slow-growing taxa. Using data from bacterial and metazoan systems, we present empirical evidence that the developmental order of ecological systems moves along the paths of least environmental resistance. The capacity of simple principles to explain the trend in the developmental order of diverse ecological systems paves the way to an enhanced understanding of the collective features characterizing the diversity of life.

## Introduction

One of the most iconic thought experiments in the life sciences is what would happen if the tape of life on Earth were rewound and played again [1]. Would the same species return in the same or similar order? Or do historical accidents lead to unpredictable outcomes? This question is not just philosophical or counterfactual, because developmental processes (the appearance of a group of taxa) undergone by complex living systems are continuously replicating the experiment [2]. Ecological systems ranging from microbes to forests are formed by self-organizing networks of interacting taxa, whose emergent behavior is crucially influenced by their history, changing biotic and abiotic environment, and chance [3]. From the primary development of microbiota in newborns to the secondary development of macrobiota in grasslands due to environmental perturbations or seasonal patterns, local tapes of life are constantly starting or being rewound. Such development over ecological timescales is analogous to evolutionary outcomes derived from similar developmental processes, increasing the relevance to understand general basic processes [4–7].

The ubiquitous collective pattern of orderly and sequenced development of ecological systems (also known as succession [2]) is widely considered to be one of the foundational problems of ecology [8–10], and it remains central to contemporary research on biodiversity and environmental change [11–13]. Many mechanisms and trade-offs promoting development have been documented, predominantly in plants [2, 14, 15]; but these emerge at the interface of specific organismal traits (e.g., dispersal ability, resource requirements, relative growth rates) and local systemic contexts (e.g., environmental stressors, prevalence of facilitative vs. inhibitory interactions). As a result, fundamental questions remain unanswered about the generality of these mechanisms and the inevitability of collective patterns or particular developmental pathways [16–18]. A generalized theoretical framework capable of positing testable predictions about developmental trajectories across the tree of life, with minimal assumptions about specific organismal traits and environmental contexts, would aid in answering these questions and advance the understanding of one of ecology’s oldest and still-paramount challenges [19–21].

One key insight is that the developmental order displayed by ecological systems is generally a process from fast-growing to slow-growing taxa [2, 22, 23]. Yet, it remains unclear whether this collective pattern depends critically on the complex combination of specific life-history traits and trade-offs that are often strongly linked to growth rate (e.g., dispersal ability) or whether it might simply be the result of local taxon interactions following the path with the least environmental resistance for species survival (requiring less adaptations), and hence the configuration upon which systems will probabilistically converge [24–26]. If there is path convergence, it becomes critical to understand the role of individual traits in modulating such local taxon interactions and paths. Indeed, discovering minimal processes that can explain the development of diverse ecological systems can shed new light on the conditions under which the tape of life might have a relatively predictable sequence of events [10, 27]. This is increasingly a practical concern, too, as conservationists seek to restore functional biodiversity in areas where species have been driven locally extinct [28–30] and as the essential roles of microbiome development in sustaining both human and environmental health become clearer [31].

Formally, we define development as the self-organized process of moving from an empty set (a novel environment with no taxa) to a community of *n* taxa (a modified environment with a particular set of taxa) through a collection of different taxon subcommunities. Previous work in experimental and theoretical ecology has established foundational knowledge linking community composition with historical events, assembly and disassembly dynamics, and invasibility [32–41]. This body of work has assumed that the environments remain relatively constant throughout the development of ecological systems (i.e., no chance events) [42]. However, the survival of species depends on historical accidents as well as on their capacity to modify and withstand changes in the abiotic and biotic environments (e.g., resource availability, competition, temperature variation) [43]. At intermediate levels of biological organization [44], the possible solutions to these environmental challenges can be represented by the possible sets of interacting populations (i.e., communities) that partition or modify the resources available at a given place and time [23, 43]. Consequently, it becomes imperative not only to understand which environments give rise to developmental processes, but also which constructed environments enabled the persistence of such processes [6, 45].

Here, we focus on the developmental order of ecological systems (i.e., knowing which taxa should be expected to appear before others regardless of whether the former taxa can survive across the entire developmental process). That is, we are not focusing on the actual step-by-step assembly and disassembly of an ecological system, but simply on the expected order of taxon appearance. Specifically, our research goal is to investigate the extent to which a minimal model rooted in local taxon interactions, but without invoking the complexity of multiple life-history traits, can explain the developmental order from fast-growing to slow-growing taxa generally observed in nature.

### Theoretical framework

We operationalize development using a general Markov chain process moving from an empty set ∅ to a given set of *n* taxa (Fig. 1; Sec. S6). Conceptually, we assume that development is possible by the compatibility (feasibility) of environmental conditions with current and future taxon communities. In our theory, development takes the path of least environmental resistance (i.e., higher compatibility of environmental conditions across a sequence of taxon communities). As a metaphor, one can think about the compatibility of environmental conditions with current and future communities as the flow of water between reservoirs. The thicker the pipeline between two adjacent reservoirs, the faster the water flows (the higher the compatibility). The thickness of a pipeline is a function of the water capacity of such reservoirs. The water capacity is constrained by the nature of building elements and the interactions between these elements. Paths (sequence of pipelines) with higher thickness represent lower environmental resistance. Water would tend to flow through the thickest pipelines imposing less friction (Figs. 1D–1E). The probability of reaching a target reservoir from an initial reservoir after *x* time steps is given by the product of probabilities (proportional to the thickness of pipelines) of using all possible path combinations across *x* pipelines. In the case of a simultaneous developmental process, there is just one pipeline. We explore how the probability of reaching a target reservoir changes under sequential and simultaneous processes (Fig. 1F).

**Figure 1:**
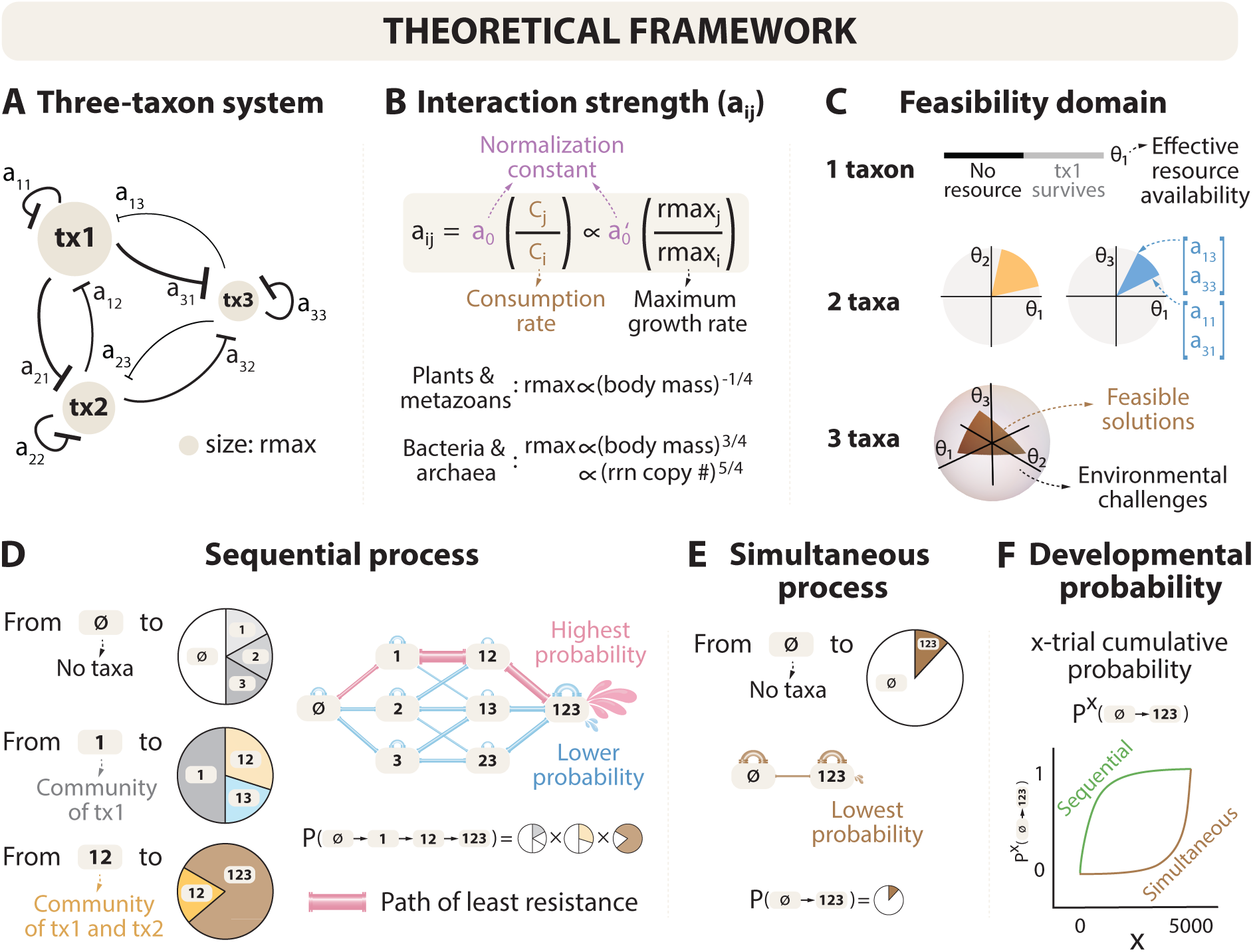
An illustration of developmental processes in ecological systems. (A) A three-taxon competition system (taxon pool: tx1, tx2, and tx3), where node size represents the maximum growth rates (rmax) of each taxon. The thickness of arrows represents the strength of pairwise interactions. (B) Estimation of pairwise interactions *a_ij_* based on rmax. Following metabolic scaling theory, rmax scales with body mass in plants and metazoans with a power of *−*1*/*4 and in bacteria and archaea with a power of 3*/*4. For bacteria and archaea, rmax also scales with the copy number of ribosomal RNA operons (rrn) with a power of 5*/*4 (Secs. S3, S5, and S10) (C) Feasibility domains for communities that consist of one taxon (e.g., community of tx1 in grey), two taxa (e.g., systems of tx1 and tx2 in yellow, or tx1 and tx3 in blue), and three taxa (in brown). The feasibility domain of a single taxon is assumed to be 0.5 (when its effective resource availability *θ*_1_ *>* 0). For a multi-taxon community, the feasibility domain is governed by the interaction strengths *a_ij_* between taxa (Sec. S2). (D) Sequential development can be probabilistically described by a Markov chain, where the feasibility-based transition probabilities *P* are shown as pie charts for communities starting from none ∅ to all three taxa *{*123*}*. These transition probabilities are determined by the feasibility of all possible communities in the sequence (match with colors in (C); Sec. S6). Each community is assumed to either maintain the current taxon composition or transition to a larger community with an additional taxon. The results remain robust considering disassembly processes (Sec. S9). The overall probability of developing the three-taxon system along the path ∅ *→ {*1*} → {*12*} → {*123*}* can be calculated by multiplying the three transition probabilities along the path. The path with the highest probability is called the path of least resistance (represented by pink water pipes). (E) Simultaneous development can also be probabilistically described by a Markov chain and is equivalent to a binomial process (Sec. S7). The transition probabilities are determined by the feasibility of *{*123*}* only (match with colors in (C); S6). (F) When comparing the probability of developing from the empty set ∅ (standardized baseline) to the largest community *{*123*}* with all taxa, both sequential and simultaneous processes are guaranteed to succeed (*y* = 1) after a certain number of trials (*x* -axis). However, sequential development can achieve this with fewer trials.

The environmental resistance of developing to a new possible community (conditional probability of moving to state *j* given current state *i*) is a function of the feasibility of such communities (Fig. 1C). The higher the feasibility of the current community and the lower the feasibility of the future community, the higher the environmental resistance of developing to a new community (see Sec. S2 for a formal definition of feasibility). Feasibility is a function of local taxon interactions (Figs. 1A–1B) and is formalized following five key assumptions: (1) All taxa have equal dispersal rates. This means that the order of appearance is unbiased but this order can be stochastic. (2) Taxon interactions are a function of taxon consumption rates. Taxa with higher consumption rates have higher competitive effects on other taxa (Fig. S20). (3) All taxa have the same carrying capacity. This means that the higher the per capita growth rate of a taxon, the higher its consumption rate and consequently the higher its competitive advantage over available resources. (4) For modeling purposes, and for application to particular data sets, the maximum growth rate of a taxon is proportional to its average body size using metabolic scaling laws. (5) The average body size is not expected to change over time. This implies that competitive effects and the feasibility of a community are also time-invariant. We present the formal feasibility analysis in Sec. S1, which combines ecological dynamics, metabolic scaling theory, and Markov processes (Fig. 1).

Using our proposed framework, we theoretically investigate the probability of a sequenced developmental process (introducing one taxon at a time) as opposed to a simultaneous developmental process (introducing all taxa at the same time). We then evaluate the conditions under which the sequential developmental process can follow an orderly path from fast-growing to slow-growing taxa. Specifically, we address whether this orderly path corresponds to the theoretical path with the highest probability, referred to as the path of least resistance. We focus our analysis on the special case of directional assembly (once a taxon is successfully introduced it cannot go extinct). Effectively, this analysis aims to explain the probability that a group of *n* species can be found together at a given point in time. While this analysis is a simplification of the developmental process undergone by natural systems (i.e., no extinctions); under our assumptions, this analysis generates the same results as if additionally considering disassembly processes (Figs. S3 and S4; Sec. S9). That is, under assembly only, the question is which path is the system most likely to take. Under assembly and disassembly, the question would be which path is most frequently visited by the system if a system is observed with *n* taxa. We test our theoretical predictions using empirical developmental data across a wide range of taxonomic levels including infant gut and marine microbiota [46, 47], tree and bird systems [48, 49], and multiyear and seasonal mammalian herbivore systems [50, 51]. Using these empirical data, we investigate the extent to which the observed developmental order follows the theoretical path of least resistance.

## Results

### Theoretical predictions

We started our theoretical analysis by investigating whether a sequenced assembly (a special case of development) is more likely than having all taxa introduced at once. In other words, is there an inherent tendency towards sequenced assemblies even without invoking extensive details about individual characteristics (only local taxon interactions)? Is development (and ecological restoration) more likely to succeed when proceeding one species at a time rather than in batches? To test these ideas, we calculated the probabilities for sequential and simultaneous development using model-generated systems following our proposed feasibility analysis (Secs. S6 and S7).

Figure 2A shows the cumulative probabilities of reaching full development as a function of the number of trials, comparing sequenced versus simultaneous development in a hypothetical system of *n* = 6 taxa, starting from different initial numbers of taxa (1 *≤ m ≤ n*). Note that *m* = *n* is equivalent to the simultaneous development (i.e., all taxa appear simultaneously). Results are qualitatively consistent using other dimensions *n* (Fig. S2). By construction, these systems can be assembled starting from any initial community (Sec. S6). We found that the smaller the initial number of taxa (after the empty set ∅), the larger the cumulative probability of development. Only after an extremely large number of attempts does the cumulative probability of simultaneous development asymptotically converge to one. Together, these results suggest that, barring a vast number of introduction attempts, it is far less likely to observe a system developed from the simultaneous appearance of many taxa than one arising from a sequenced process.

**Figure 2:**
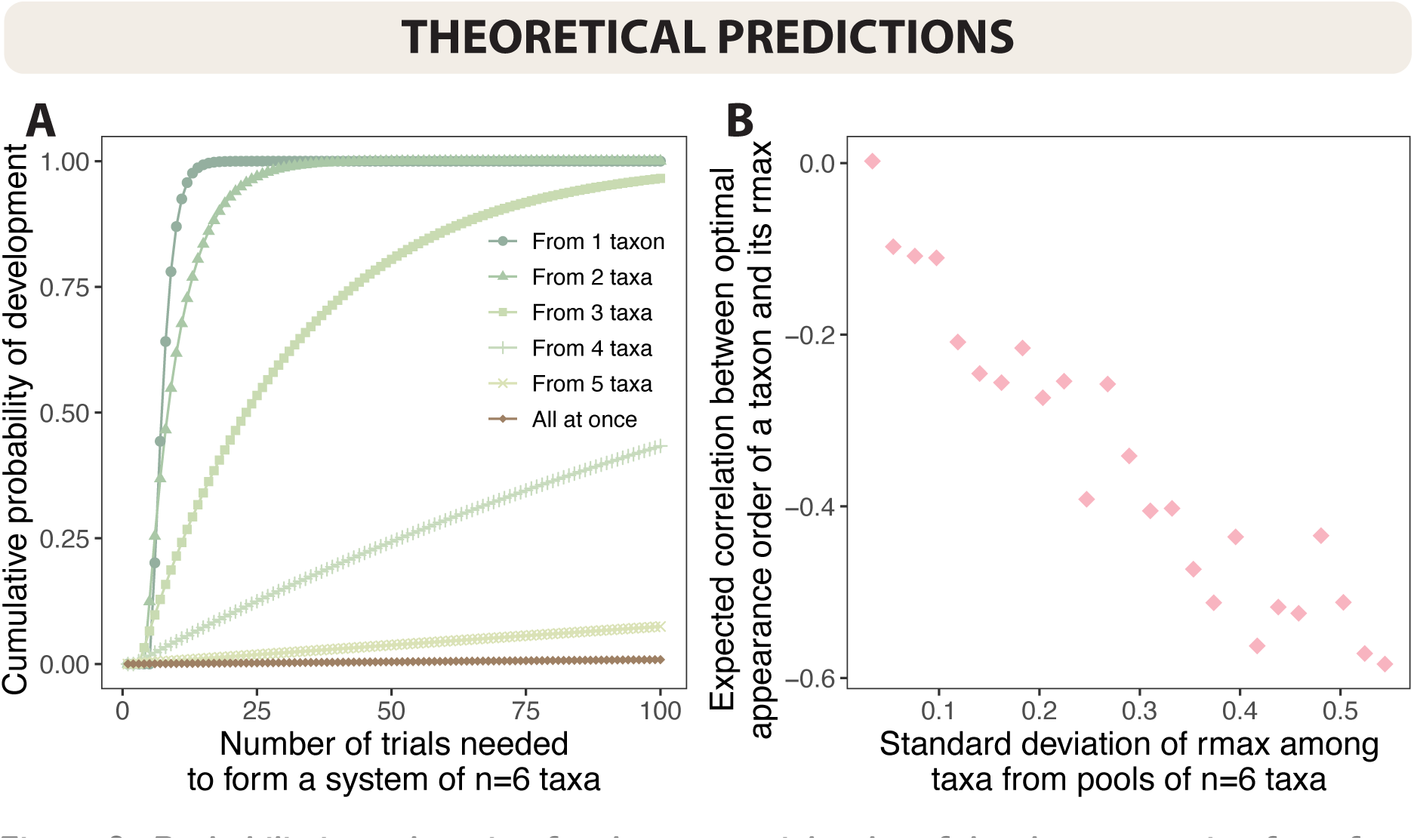
Probabilistic explanation for the sequential order of development going from fast-growing to slow-growing taxa. (A) Comparison of cumulative probability of development (*y* -axis) between sequential development (green) with varying numbers of initial taxa and a simultaneous development (brown) within a limited number of trials (or attempts; *x* -axis). Notably, in sequential development, the process starting from one taxon reaches probability 1 more quickly compared to those starting with larger initial communities. Additionally, the probability of development through the simultaneous process is negligibly small, though it may reach probability 1 if the number of attempts is sufficiently large (Secs. S6 and S7). (B) Relationships between the path of least resistance and rmax of sequential development starting from 1 taxon. Firstly, the negative expected correlations between the most likely developmental order of a taxon and its rmax (*y <* 0; Spearman’s rank correlation) indicate that, in order to maximize the probability of development, fast-growing taxa with larger rmax are expected to appear earlier. Also, as the standard deviation of rmax among taxa (*x* -axis) increases, these negative correlations tend to be stronger (approaching *y* = *−*1). This finding suggests that when the diversity in the rmax among taxa within a system is higher, the appearance order starting from fast-growing taxa to slow-growing taxa becomes more distinguishable. Note that in both panels, we choose *n* = 6 taxa only for illustrative purposes. The results remain consistent across different choices of *n* (Fig. S2) and when additionally considering disassembly (Figs. S3 and S4; Sec. S9).

Having established the inherent probability of a sequenced development of ecological systems, we theoretically investigated the conditions under which it follows a sequenced orderly process as a function of biodiversity. To do so, we used the same systems of *n* = 6 taxa generated before (as an illustrative example). Then, we divided these model-generated systems into groups, ensuring equal intervals of the standard deviations of rmax among taxa. For each system, we identified the path with highest probability, that is the path of least resistance. Next, we calculated Spearman’s rank correlation between the taxon order in each estimated path and the generated taxon rmax. Figure 2B shows that the most likely appearance order of a taxon is, in general, strongly and negatively correlated with its rmax, indicating that the path of least resistance occurs when fast-growing taxa appear first. However, note that positive correlations between appearance and rmax can be observed, especially in unlikely paths. Furthermore, we found that higher variations in growth rates are associated with more strongly negative correlations between a taxon’s most likely appearance order and its rmax. By extension, pools with a greater diversity of taxa accentuate the positive association between developmental success and a successional process going from fast-to slow-growing taxa.

Because our feasibility analysis is based on local taxon interactions, our results can be explained by the fact that similar competitive advantages (similar rmax) promote symmetric interactions, and in turn increase the feasibility of a community (Fig. S20). Asymmetries get amplified between taxa with small competitive advantages (Fig. S21). Similarly, because the probability of a path is given by the multiplication of conditional probabilities, this product is maximized by having taxa with similar rmax together in an ordered assembly rather than a disorderly one. Theoretically, the path of least resistance solves a Pareto front maximizing the probability of staying and leaving. For example, this is achieved at *P*(staying) = *P*(leaving) = 0.5. This means that the path cannot select the maximum probability at each step, only the maximum product of probabilities, implying a combination of forming and breaking symmetries. By starting from fast-growing taxa, the probability of developing the system (the largest community) under a Markov process increases, but it is difficult to find a perfectly ordered process unless there is a high variability of rmax among taxa. While all these previous theoretical results are generated assuming a directional assembly (i.e., no extinctions); under our assumptions, these results do not change by following more complex scenarios involving disassembly dynamics (Figs. S3 and S4; Sec. S9). These results imply that biodiversity can turn into a heterogeneity of local taxon interactions, which can then amplify the emergence of a sequenced and orderly development.

### Empirical corroboration

To empirically evaluate our theoretical predictions, we compiled publicly available data from developmental processes involving taxonomic levels ranging from species to phyla, timescales spanning days to decades, and organisms spanning *∼*15 orders of magnitude in size from bacteria to elephants (Sec. S11). The first two datasets represent primary development in microbes: the development of an infant gut microbiota comprising six main phyla across 2.5 years using fecal samples and 16S rRNA and metagenomic sequence data [46], and the development of marine bacteria from a laboratory model system comprising five main orders over a span of 144 hours [47]. The third and fourth datasets represent secondary development in trees and birds: the development of a community of trees comprising 7 species in Princeton, NJ, that was disturbed by a blight in 1920 and hurricanes in 1938 and 1944 [48], and the development of an assemblage of birds comprising 13 main families from abandoned agricultural fields [49]. The fifth and sixth datasets represent secondary and seasonal development, respectively, of large mammalian herbivores in Gorongosa National Park: the development of the entire herbivore system comprising 16 genera over 28 years after the Mozambican Civil War [51], and the development of a local herbivore system comprising 11 genera throughout the dry season in 2016 [50]. Maximum growth rates (rmax) were approximated using metabolic scaling relationships based on average ribosomal RNA operons copy numbers for bacteria and archaea [52], diameter at breast height for trees [53], and body masses for metazoans [50, 54].

Figure 3 shows that the development of these empirical systems tends to proceed from fast-growing to slow-growing taxa. As predicted, we found strong negative correlations (Spearman’s rank correlation *ρ ∈* [*−*0.93, *−*0.44]) between the observed order of taxon appearance and the rmax estimated for each taxon. Next, for each empirical system, we determined the theoretical path of least resistance using rmax following our proposed feasibility analysis (Sec. S8). In line with our theoretical results (Fig. 2), we found strong positive correlations (Spearman’s rank correlation *ρ ∈* [0.48, 0.93]; pink stars) between the observed order of taxon appearance and the theoretical optimum, which were always larger than those expected from random appearance orders (boxplots). These results are robust to various assumptions and criteria used in data processing (Figs. S5 and S6; Sec. S11).

**Figure 3:**
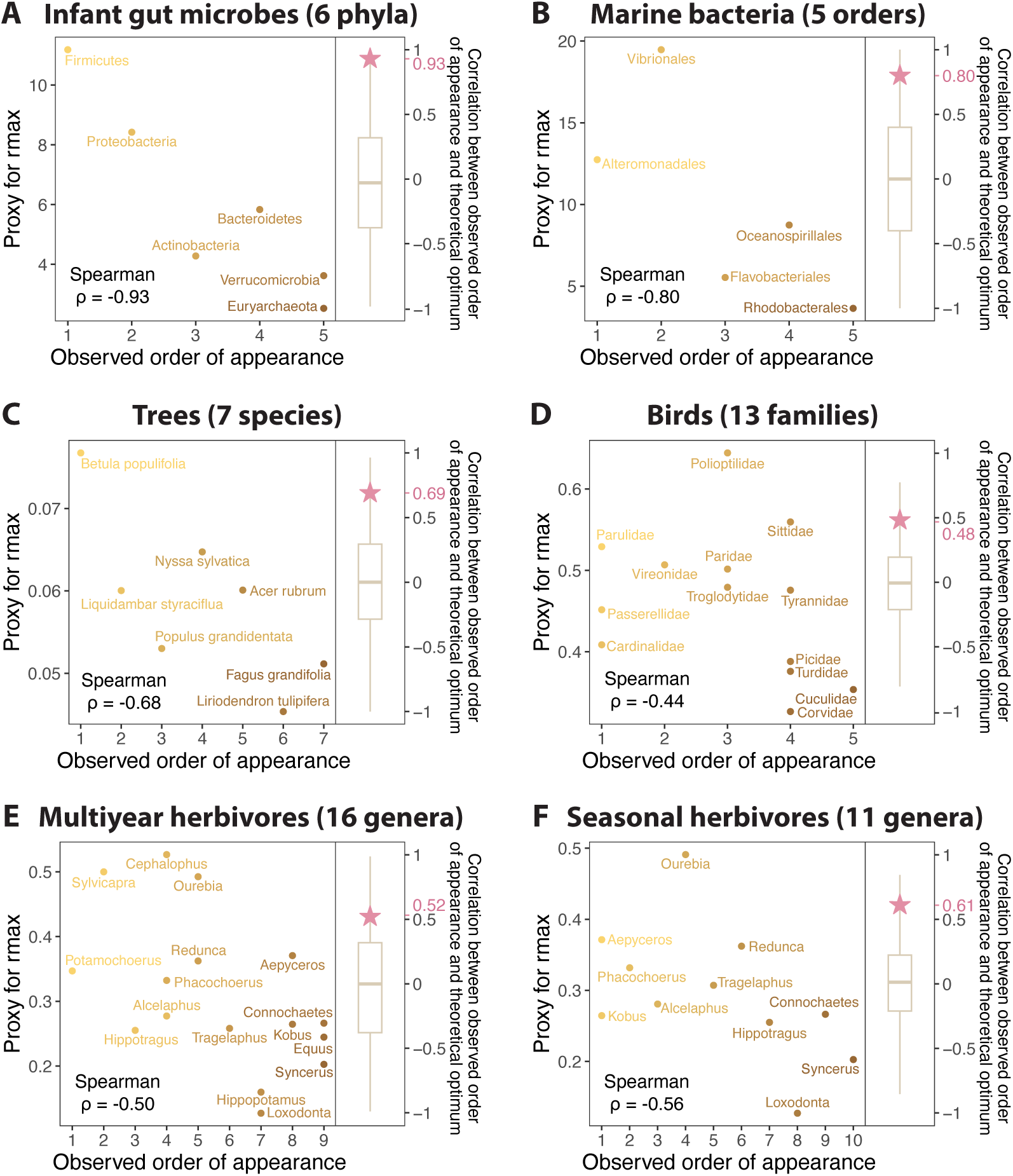
Observed developmental order in empirical systems and comparisons with theoretical and random orders. (A) Primary development involving 6 phyla of gut microbes in a healthy male infant [46]. (B) Primary development involving 5 orders of marine bacteria in a laboratory study [47]. (C) Secondary development involving 7 species of trees in the northeastern United States [48]. (D) Secondary development involving 13 families of birds in the southeastern United States [49]. (E) Secondary development (corresponding to the postwar reassembly over the period 1994–2022) involving 16 genera of herbivores in Mozambique [51]. (F) Seasonal development (over the dry season in 2016) involving 11 genera of herbivores in Mozambique [50]. In each of these empirical systems, the correlation between the observed order of appearance (*x* -axis) and the proxy for rmax (*y* -axis) is negative (Spearman’s rank correlation *ρ* = *−*0.93, *−*0.80, *−*0.68, *−*0.44, *−*0.50, and *−*0.56 in (A)– (F), respectively), in line with our theoretical prediction (Fig. 2B). On the right side of each panel, the correlation between the observed order of appearance and the theoretical order of path of least resistance (pink stars; *ρ* = 0.93, 0.80, 0.69, 0.48, 0.52, and 0.61, respectively) is significantly higher than expected under randomly generated orders (boxplots; based on 10^3^ random paths). Please refer to Sec. S11 for more details on each empirical system.

Additionally, Figure 4 shows that development from fast-to slow-growing taxa becomes stronger with increasing diversity (variation) in their growth rates, confirming our theoretical predictions. For each empirical system, we drew all possible subsets (combinations) consisting of 60% taxa. Then, for each subset, we calculated Spearman’s rank correlation between the observed order of appearance of a taxon and its rmax, as well as the standard deviation of rmax among all taxa within the subset. Next, we divided all subsets into three diversity groups based on the standard deviations of rmax and plotted the distribution of correlations between observed appearance order and rmax within each group. In line with our theoretical predictions (Fig. 2B), we found that average correlations become more negative with greater diversity in rmax. These results are robust to the choice of taxon pool size and the number of diversity groups (Figs. S7–S18), suggesting that diversity plays a central role in shaping the development of ecological systems [55].

**Figure 4:**
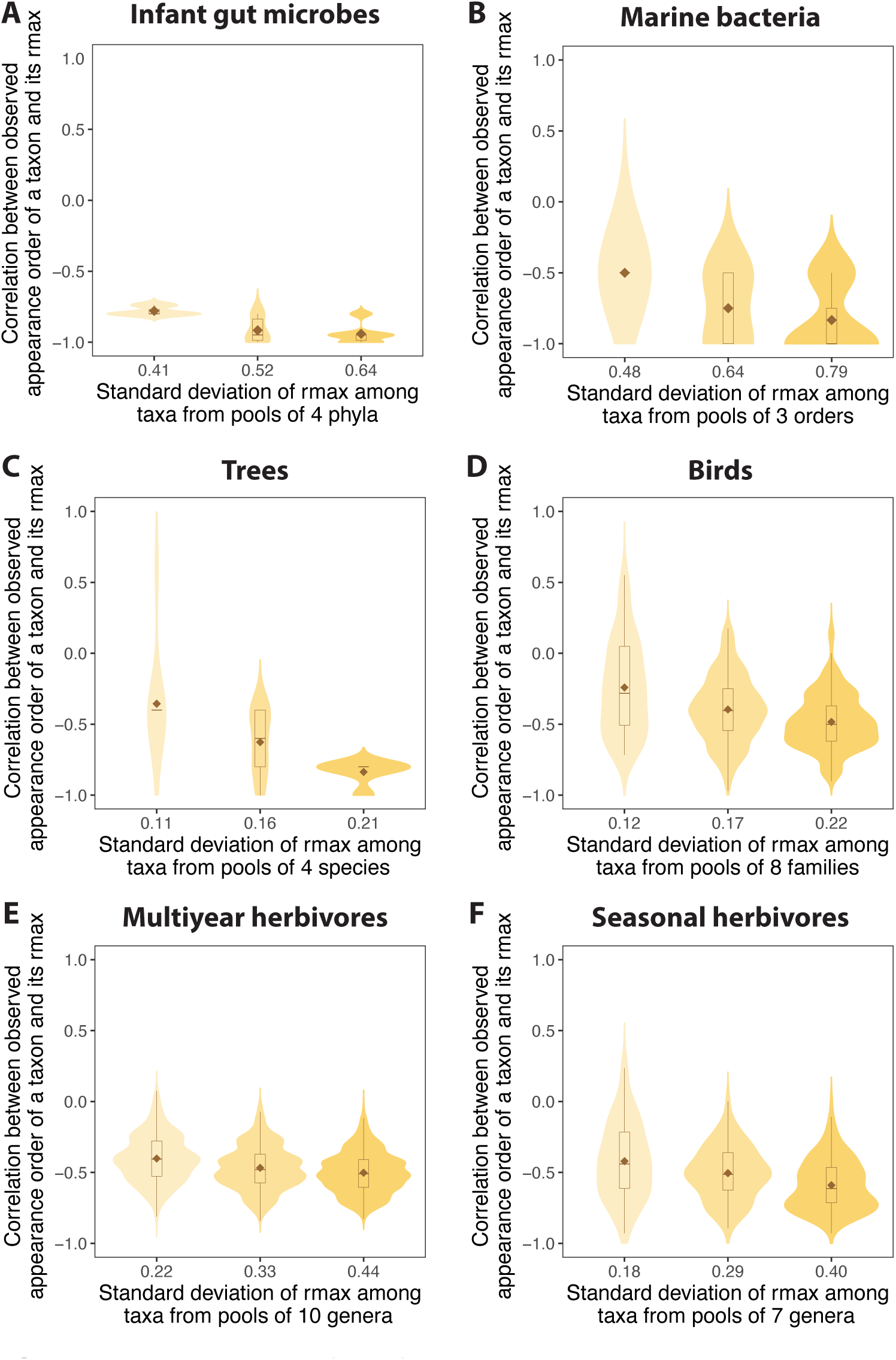
Sequential development from fast-growing to slow-growing taxa becomes more distinguishable as the diversity in rmax among taxa increases. (A)–(F) For each of the empirical systems in Fig. 3, violin plots and embedded boxplots show the distribution of correlations between the observed appearance order of a taxon and its rmax (*y* -axis; Spearman’s rank correlation) as a function of the standard deviation of rmax among taxa (*x* -axis). Brown diamonds inside each boxplot show the expected value (mean) of the correlations. As the standard deviation of rmax increases, the correlation between the observed order of appearance and rmax becomes stronger (approaching *y* = *−*1), in line with our theoretical prediction in Fig. 2B. Here, we choose the pool size as 60% of the entire system size and set the number of diversity groups to three, but results are consistent across different choices of parameters (see Figs. S7–S18).

## Conclusions

We have presented a general theoretical framework for development based on basic feasibility assumptions rooted in local taxon interactions to better understand the conditions underlying the tendency of taxa to appear from fast-growing to slow-growing taxa. More than a century of evidence shows that the development of ecological systems is characterized by ordered processes, and that systems often converge on similar trajectories [2, 11, 14, 56]. Can this phenomenon be explained only with reference to a large combination of individual traits and circumstances, or can it be understood as a self-organized collective outcome emerging from basic premises? We have shown that if one assumes a given developmental order as a local solution via taxon interactions to changing environmental challenges, then the solution that maximizes the number of surmountable environmental challenges is typically the one that proceeds from fast-growing to slow-growing taxa. In other words, development follows paths of least environmental resistance. Because biodiversity promotes an orderly sequential development and natural selection acts at the local level [44], it can be speculated that eco-evolutionary dynamics via life-history traits and environmental compatibility may play a reinforcement role in preserving this basic developmental process at the system level [6].

Supporting the statements above, we have shown theoretically that a sequential development is more likely than a simultaneous development, at least on the scales of time and iteration that plausibly resemble those of natural events. Indeed, it requires an enormous amount of tape repetitions (attempts) for a simultaneous assembly to become comparable to a sequential one. Another prediction derived from our theory is that the higher the heterogeneity of intrinsic growth rates within a pool of taxa, the more pronounced the ability of an orderly process can characterize the development of ecological systems—perhaps helping to explain why changes in biodiversity can drastically alter the development of ecological systems [55]. Importantly, we have focused on paths that can minimize environmental resistance and become more likely. While there is a tendency in these paths to go from fast-growing to slow-growing taxa, small differences in growth rates can introduce deviations from this development structure. For example, variations in life-history traits or life-history trajectories can help to explain deviations from the expectation [57]. It is worth mentioning that these results do not provide the specific step-by-step order of assembly (and disassembly), but simply they provide a minimal model rooted in local taxon interactions to explain the expected order of taxon appearance in natural systems without the need to invoke a high level of information about life-history traits. That being said, this minimal model implies the potential existence of general rules or principles that can apply to the development of life on Earth.

Using empirical data from a wide range of developmental processes and taxonomic groups, we have found support for our theoretical results and predictions. The qualitative consistency of these results from very small ecological systems (bacteria in 1-litre bottles under controlled laboratory conditions over 6 days) to some of the world’s largest (ungulates and elephants in a 4,000 km^2^ national park) suggests a potentially high level of generality. We emphasize, however, that our results do not rule out alternative hypotheses of developmental processes, nor are they incompatible with explanations that invoke more nuanced biological mechanisms to explain specific trajectories. Rather, our results illustrate that basic assumptions can be sufficient to explain coarse-grained properties of these dynamics and establish an expectation for their common tendencies. Future studies might use our work as a platform to explore general paths and processes for taxonomic and interaction turnover during development. Indeed, the interplay between stochasticity and determinism continues to be an active line of research [13, 16, 58–60]. On the one hand, historical contingency (environmental conditions) seems to be an underlying factor modulating the short-term dynamics of ecological systems [1, 23]. On the other hand, convergent evolution seems a plausible explanation for finding a finite set of long-term solutions to an environmental challenge [56, 61]. Integrating these two views, the question is whether these potential finite solutions are themselves compatible with a collection of different environmental conditions, where the larger the collection, the larger the feasibility of observing a solution [62, 63]. Our work suggests that accounting for this feasible least resistance will be a step towards the longstanding aim of understanding the expected outcomes of repeating the tape of life.

### Data and code availability

This paper analyzes existing, publicly available data (see Section S11). All original code has been deposited at Github: https://github.com/MITEcology/Deng_Assembly. Any additional information required to reanalyze the data reported in this paper is available upon request.

## Acknowledgments

J.D. and S.S. thank the financial support from MathWorks. R.S. thanks the support of the Generalitat de Catalunya under grant AGAUR 2021 SGR 00751. R.M.P. thanks the National Science Foundation (BoCP DEB-2225088), the Carr and Cameron Schrier Foundations, the Biodiversity Grand Challenge of the High Meadows Environmental Institute, and Gorongosa National Park’s Department of Scientific Services. S.A.L. thanks NSFDMS 1951358. The authors thank the contributors and maintainers of databases rrnDB, BIEN, and AVONET for their valuable work. Furthermore, the authors acknowledge the MIT SuperCloud and Lincoln Laboratory Supercomputing Center for providing High-Performance Computing resources that have contributed to the research results reported within this paper.

## Author contributions

Conceptualization, all authors; Methodology, J.D., R.M.P., S.S.; Software, J.D.; Validation, all authors; Formal Analysis, J.D., R.M.P., S.S.; Investigation, all authors; Resources, O.C., R.M.P.; Data Curation, J.D.; Writing – Original Draft, J.D., R.M.P., S.S; Writing – Review & Editing, all authors; Supervision, S.S.

## Supplementary Materials

### S1 Feasibility analysis

We illustrate the proposed feasibility premises of development using a hypothetical, competition, three-taxon system *S* as an example (Fig. 1). We assume that the partition or modification of resources is characterized by the competitive effect *a_ij_* = *a*_0_(*C_j_/C_i_*) of taxon *j* on taxon *i* (Figs. 1A–1B), where *a*_0_ is a normalization constant and *C_i_* is the average consumption rate of taxon *i* (Eq. (S15); Sec. S3). To study the maximum effect of growth rates on competition, we assume that all taxa have the same environmental carrying capacity and dispersal ability, and consequently *a_ij_ ∝ a*_0_*^′^* (rmax*_j_/*rmax*_i_*), where rmax*_i_* is the maximum growth rate of taxon *i* (Eq. (S35); Sec. S5). This assumption implies that taxa are not constrained to enter at a particular order based on their specific traits (e.g., dispersal ability) other than their maximum growth rates and the current subset of present taxa [1–3]. Next, using metabolic scaling theory [4], we can approximate maximum growth rates as a function of average body mass (*M_i_*) as rmax*_i_ ∝ M^β^*, with *β* = *−*1*/*4 for plants and metazoans, and *β* = 3*/*4 for bacteria and archaea [5, 6]. For bacteria and archaea, maximum growth rates can also be approximated as a scaling function of ribosomal RNA operons (rrn) copy numbers as rmax*_i_∝ rrn^β^* with *β* = 5*/*4 [7] (Fig. 1B). The choice of scaling exponent does not affect the qualitative results of our framework (see Sec. S10). It is worth noting that our proposed framework can be considered as a basic process upon which one can add more complex contexts. For example, our framework illustrates that carrying capacities can act as an upper limit to consumption rates and can be used instead of growth rates upon the selection of specific trade-offs (Eq. (S28); Fig. S19; Sec. S4).

We also assume that any possible subset of taxa (community *C*) from the hypothetical system (*C ⊆ S*) is compatible with a specific *feasibility domain* of environmental space *D_F_* (*C*), and its feasibility is proportional to the size of such a domain and is a function of taxon interactions (Fig. 1C). Formally, we determine *D_F_* (*C*) following the Lotka-Volterra model (Eq. (S1)): d*N_i_ /*d*t* = *N_i_ r_i_ /θ_i_ θ_i_ − _j∈C_ a_ij_ N_j_*, where *N_i_* is the density of taxon *i*, *r_i_* is the intrinsic growth rates of taxon *i*, *θ_i_* is the effective resource availability of taxon *i*, and *a_ij_* is the competitive effect described in the preceding paragraph. Then, the feasibility domain corresponds to the collection of compatible environmental conditions (*D_F_* (*C*) = *_j∈C_ N_j_^∗^*a*_j_* : A = [a_1_ a_2_ *· · ·* a*_|C|_*], *N_j_^∗^ >* 0 *∈* R*^|C|^*) leading to positive equilibrium densities (N*^∗^* = A*^−^*^1^*θ >* 0) constrained by the column vectors (a*_j_*) of the competition matrix A (Sec. S2). Because we are interested in a developmental process, we assume that the system *S* can exhibit different community compositions, and therefore, a different resource partition or modification across time [1–3]. For instance, in a hypothetical system under a directional assembly, we may observe that taxon 1 appears first, then taxon 2, and lastly taxon 3. This sequence implies that changes in the feasibility domain are given by *D_F_* (*{*1*}*) *→ D_F_* (*{*12*}*) *→ D_F_* (*{*123*}*). This same logic would apply to any other possible sequence of events, representing possible solutions to time-varying environmental conditions.

Lastly, we assume that a system *S* develops through sequenced community changes (the developmental path) that have the highest probability (Fig. 1D; Sec. S8). We search for this *path of least resistance* following a Markov chain process (the solution with the highest probability). Specifically, we translate each feasibility domain *D_F_* (*C*) into a probability *P*(*C*) = vol(*D_F_* (*C*) *∩* Θ*^|C|^*)*/*vol(Θ*^|C|^*), where vol(Θ*^|C|^*) corresponds to the volume of the parameter space of the environmentally-driven parameters *θ* (a proxy for environmental space). For example,assuming that taxon 1 appears first, the f_e_asibility domain *D_F_* (*{*1*}*) corresponds to half of vol(Θ) and *P*(*{*1*}*) = 0.5. We assume *P*(*{*1*}*) = *P*(*{*2*}*) = *P*(*{*3*}*), which implies consistent dispersal probabilities across the three taxa. Because t_h_is is a competition system, *_DF_* (*{*1*}*) *> D_F_* (*{*12*}*) *> D_F_* (*{*123*}*) [8], which implies *P*(*{*1*}*) *> P*(*{*12*}*) *> P*(*{*123*}*). However, unless interactions are the same, *P*(*{*12*}*) ̸= *P*(*{*13*}*) ̸= *P*(*{*23*}*). Under a Markov chain process, a system can either remain in its current state or change (refer to sections below). In our example, the systems will transition from empty ∅ to a one-taxon community with probability 1 *− P*(∅ *→* ∅) = 0.5. But, whether it transitions to *C* = *{*1*}* instead of *C* = *{*2*}* or *C* = *{*3*}* depends on the ratio *P*(*{*1*}*) : *P*(*{*2*}*) : *P*(*{*3*}*). Assuming taxon 1 appears, the system will remain with a probability of *P*(*{*1*} → {*1*}*) = 0.5. Alternatively, it will transition to a new state with probability *P*(*{*1*} → {*12*}*)+*P*(*{*1*} → {*13*}*) = 1*−P*(*{*1*} → {*1*}*) = 0.5. But, whether it transitions to *C* = *{*12*}* instead of *C* = *{*13*}* depends on the ratio *P*(*{*12*}*) : *P*(*{*13*}*). This enables us to quantify the probability of each path by multiplying the transition probabilities along the chain (e.g., *P*(∅ *→ {*1*} → {*12*} → {*123*}*) = *P*(∅ *→ {*1*}*) *× P*(*{*1*} → {*12*}*) *× P*(*{*12*} → {*123*}*)).

Notice that the sequenced probability *P*(∅ *→ {*1*} → {*12*} → {*123*}*) is not necessarily the same as the one single event or simultaneous development *P*(∅ *→ {*123*}*) (Fig. 1E). Then, the probabilities among all possible changes can be written as a transition matrix (P) (Sec. S6). The path of least resistance is mathematically the shortest path of the directed and weighted graph constructed using the adjacency matrix *−* log P (Eq. (S44); Sec. S8).

To perform our theoretical analysis, we calculated the cumulative probabilities for sequential and simultaneous development in different model-generated systems (Fig. 1F; Sec. S6). For each system, we assign taxon body masses drawn from a log-normal distribution centered at zero with varying standard deviations. Then, we estimated taxon rmax using the generated body masses following a scaling relationship with a power of 3*/*4 (i.e., rmax = (body mass)^3*/*4^). Results are qualitatively consistent across different power relationships (Fig. S1). Next, we estimated the elements of each interaction matrix A as *a_ij_ ∝ a*_0_*^′^* (rmax*_j_/*rmax*_i_*) (Eq. (S35)), and calculated the transition matrix P for the developmental process (Eq. (S43)). The cumulative probability of development is calculated as the conditional probabilities of transitioning to a particular set of *n* taxa (community *S*) in *x* trials (attempts) starting from the empty set ∅. For each initial taxa, the respective conditional probability is the value at the intersection of row ∅ and column *S* in the *x* -step transition matrix P*^x^*. Specially, under the simultaneous development, the cumulative probability can be understood as the frequency of observing at least one success in *x* independent attempts when having all taxa together (1 *−* (1 *− P*(*{S}*))*^x^* ; Sec. S7), where *P*(*{S}*) is the one-attempt probability (Eq. (S3)). Note that the results are consistent when additionally consider disassembly (Sec. S9).

### S2 Lotka-Volterra model and feasibility

Our theoretical framework is based on the classic Lotka-Volterra model:

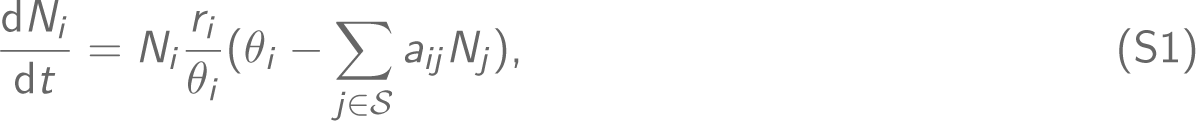

where the vector N = (*N*_1_, *· · ·*, *N_|S|_*)^⊺^ represents the densities of all taxa *i ∈ {*1, *· · ·*, *|S|} ≡ S* within the system *S*. Here, *|S|* denotes the number of taxa in the system (i.e., the cardinality of a set). The vector *r* = (*r*_1_, …, *r_|S|_*)^⊺^ *∈* R*^|S|^* represents the intrinsic growth rate of all taxa *i ∈ S*. The vector *θ* = (*θ*_1_, …, *θ_|S|_*)^⊺^ *∈* R*^|S|^* represents the effective resource availability (also known as environmental carrying capacity) of all taxa *i ∈ S*. The matrix A = (*a_ij_*) *∈* R*^|S|×|S|^* represents the pairwise interaction matrix. Each element *a_ij_* is unitless, corresponding to the per-capita effect of taxon *j* on taxon *i* relative to taxon *i*’s self-regulation. *a_ij_ >* 0, *i ̸*= *j* represent interspecific competitive interactions. Intraspecific competitive interactions *a_ii_* = 1 with the unitless normalization. Note that the classic Lotka-Volterra model Eq. (S1) is a phenomenological model. The effective resource availability *θ* = (*θ*_1_, …, *θ_|S|_*)^⊺^ *∈* R*^|S|^* consists of the joint effect of the internal (e.g., intrinsic growth rate) and density-dependence factors acting on the growth rate of each taxon [3, 9, 10].

In this paper, we assume that the pairwise interactions serve as a coarse-grained measure of the average interactions under environmental uncertainty, which is in line with a structuralist (also known as internalist) perspective [1, 11, 12]. Specifically, the structuralist perspective assumes that interactions among taxa (i.e., biological structure) are invariant and determine (or constrain) the environmental conditions compatible with the feasibility of ecological systems. Therefore, while it is expected to observe changes in pairwise interactions, the interaction matrix A can be interpreted as an expectation of the invariant biological structure of an ecological system. This conceptual simplification also allows for a rough estimation of interactions using metabolic scaling laws (Fig. 1B; Secs. S3, S5, and S10). Besides, it is worth noting that our analysis focuses exclusively on competition systems to meet the requirements of the estimation (Fig. 1A; Sec. S3).

Additionally, each effective resource availability *θ* captures the phenomenological influence of an environmental challenge on the growth potential of every individual taxon in the system *S*. This implies that distinct environmental challenges will lead to varying *θ* of the taxon. The combinations of *θ* compatible with the feasibility of taxa in the system *S* (known as the feasibility domain *D_F_*) can provide an approximation of the possibility of the system *S* in response to environmental challenges [1, 3]. Specifically, the feasibility domain *D_F_* (*S*) is bounded by the column vectors a of interaction matrix A:

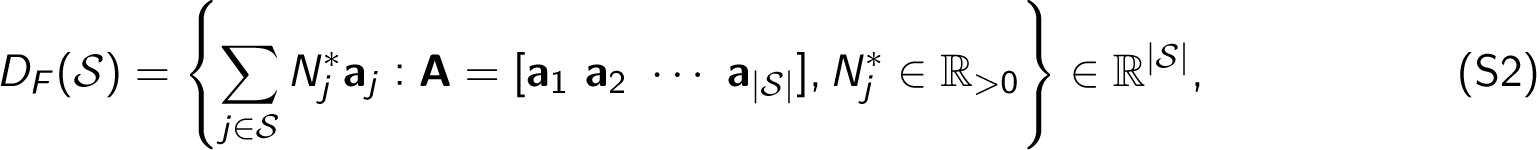

where a*_j_* is the *j*^th^ column vector of the interaction matrix A. In other words, if *θ ∈ D_F_* (*S*), it guarantees that all taxa within the system *S* are feasible (i.e., N*^∗^* = A*^−^*^1^*θ >* 0), a necessary condition for coexistence under equilibrium dynamics [13]. For any community *C* within the entire system *S* (i.e., *C ⊆ S*), the feasibility domain *D_F_* (*C*) is bounded by the column vectors of the *|C|×|C|* sub-matrix within the interaction matrix A that corresponds to the community *C*.

Without any prior information about the environmental challenges, we assume all directions of *θ* are possible and equally likely, making the parameter space Θ*^|S|^* of *θ* a closed unit sphere in *|S|*-dimensional space. Then, we can normalize combinations of compatible *θ* by the parameter space Θ*^|S|^* of *θ* to approximate the feasibility (a probability between 0 and 1; Fig. 1C):

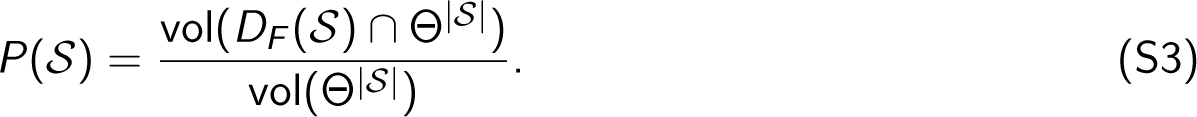

It is worth mentioning that the assumption about an unbiased environment does not affect our qualitative results. This is true because even if we bias the environment (parameter space), this biased space needs to be inside the feasibility domain of all species together (otherwise there is no assembly). This implies that on average the bias will affect all the conditional probabilities in a proportional fashion leading to nearly the same path of least resistance [3].

Furthermore, we would like to add a brief note that feasibility can be used to compute the transition probability, which represents the likelihood of transitioning from one community to another in the ecological developmental process (see Sec. S6; see also Figs. 1D–1E). Moreover, intuitively, this transition probability has the potential to indicate the most likely path of community changes (see Sec. S8).

### S3 Estimating competitive interactions (a_ij_) using consumption rates (C)

The estimation of competitive interactions *a_ij_* is derived from the Consumer-Resource equations, wherein the consumers compete for resources, thus forming a competition system. The mathematical transformation from the Consumer-Resource equations to the Lotka-Volterra (LV) competition equations refers to Reference [14].

In particular, suppose that there are two self-replicating resources, which leads to the resource equations:

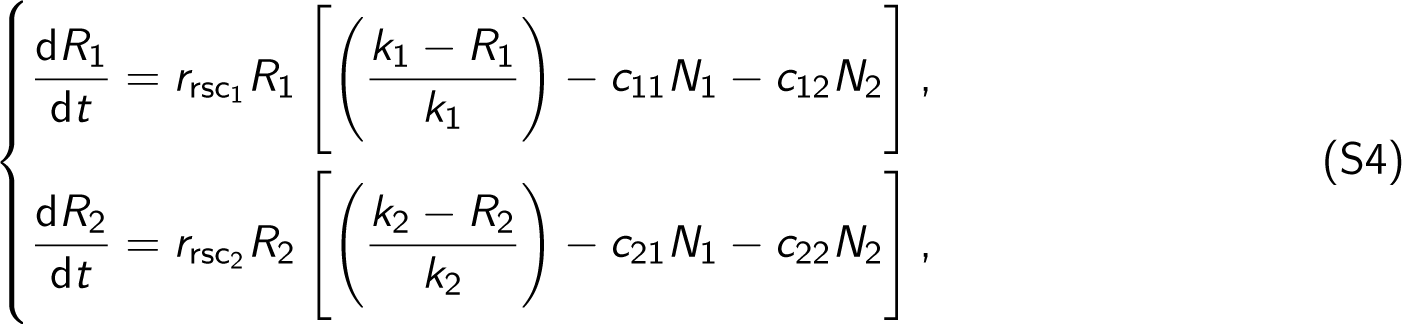

where *R_i_* is the density (or biomass) of the resource *i*, *r*_rsc*i*_ is the intrinsic rate of increase of resource *i*, *k_i_* is the carrying capacity of the resource *i*, *c_ij_* is the rate at which the resource *i* is consumed by the *j*^th^ consumer, and *N_i_* is the population density of the *i*^th^ consumer.

Then, considering two consumers, we can write the consumer equations according to the classic LV predator-prey equations [15]:

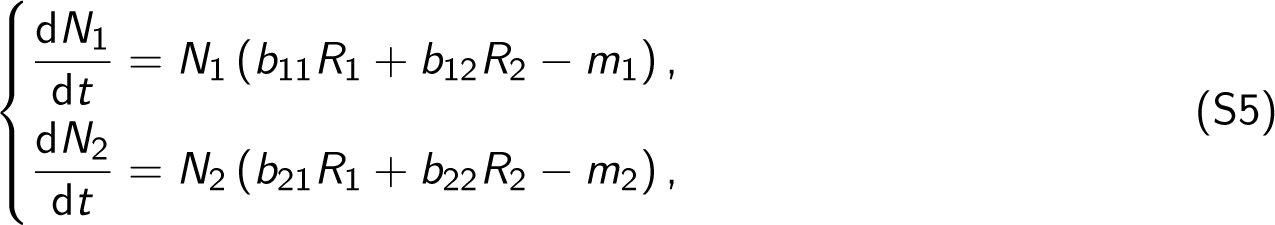

where *b_ij_* is the relative conversion of a unit of resource *j* by consumer *i*, and *m_i_* is the death rate of the *i*^th^ consumer. Note that *b_ij_* is different from mass/energy transformation efficiency.

Assuming that the dynamics of the resources occur significantly faster than those of the consumers, we can equate the derivatives of Eqs. (S4) to zero and solve for the equilibrium values *R*_1_*^∗^* and *R*_2_*^∗^* of the two resources:

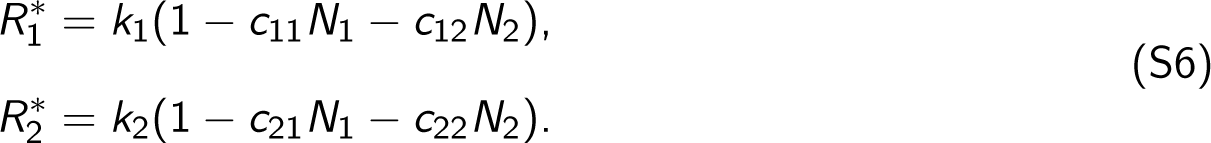

Then, we substitute *R*_1_ and *R*_2_ in Eqs. (S5) by *R*_1_*^∗^* and *R*_2_*^∗^*, respectively. That is,

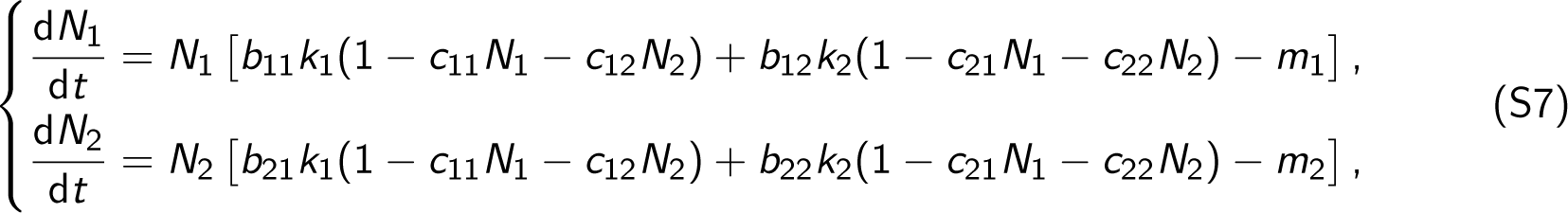

which can also be written as

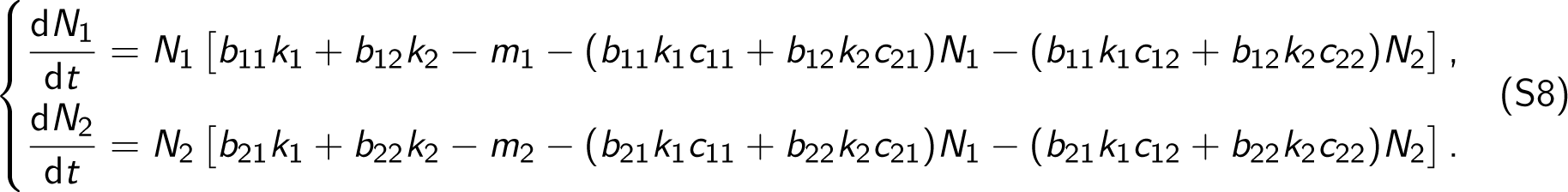

Multiplying and dividing the two equations in Eqs. (S8) by

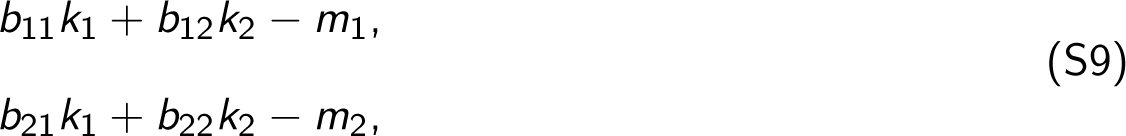

respectively, we obtain

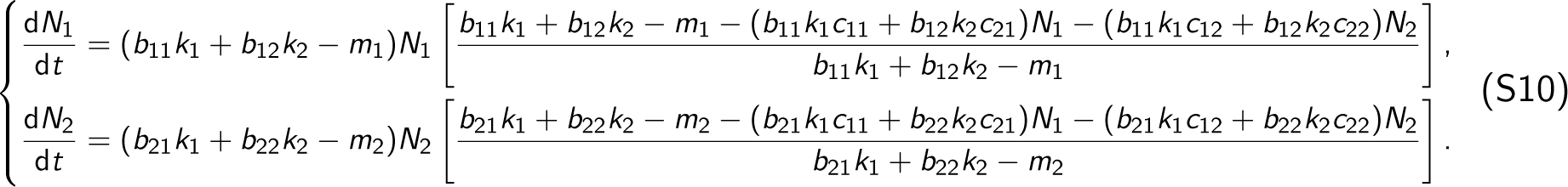

Again, dividing both the numerator and denominator of the fractions on the right sides of the two equations in Eqs. (S10) by

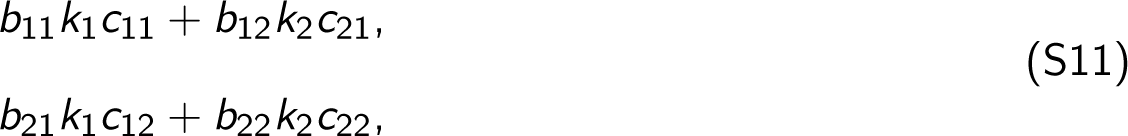

respectively, gives us

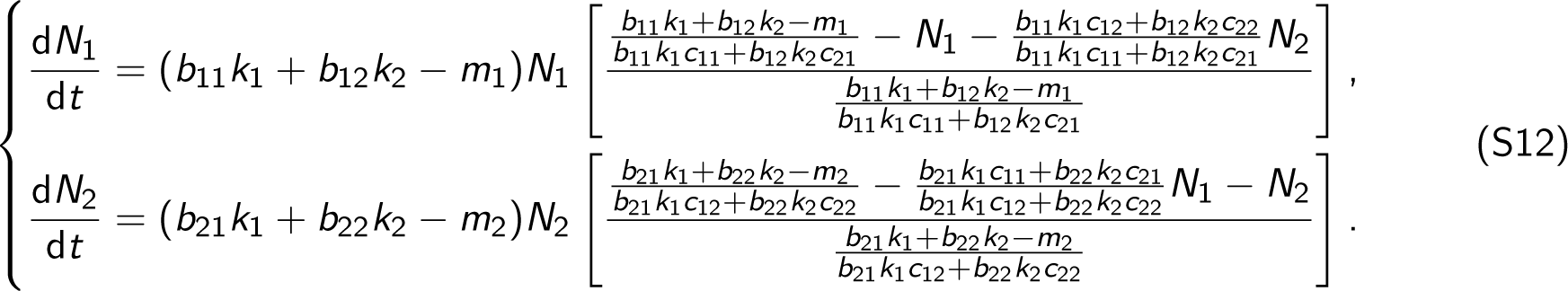

Essentially, Eqs. (S12) are the classic LV competition equations, which can be easily recognized if we make the following substitutions

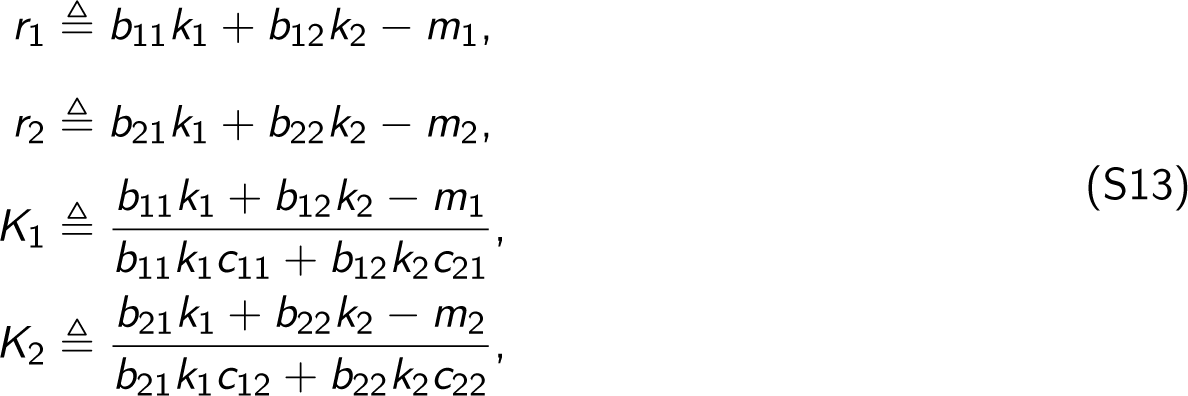

and

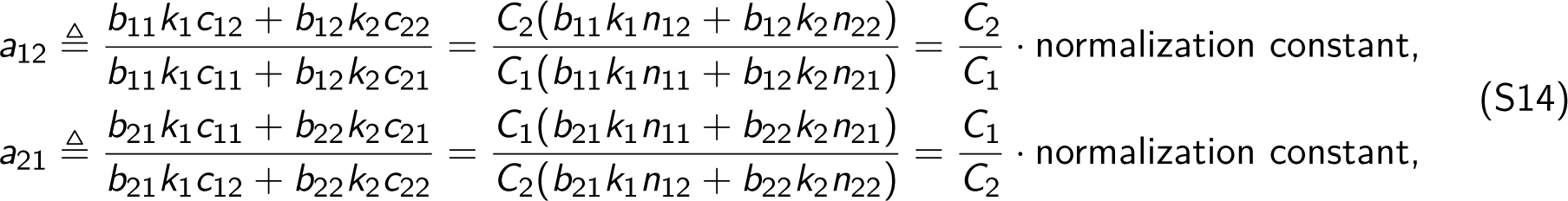

where *r_i_* is the intrinsic rate of natural increase of consumer *i*, *K_i_* is the carrying capacity of consumer *i*, and *a_ij_* is the competition coefficient measuring the competitive effect of consumer *j* on consumer *i*. Specifically, 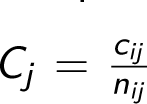 is the average rate at which consumer *i* consumes resources, where *n_ij_* is a normalization constant. Furthermore, it is typically observed that the relative conversion *b_ij_* of a unit of resource *j* by consumer *i* and the carrying capacity *k_j_* of resource *j* have limited variations for a given consumer *i* across different resources *j* and can be assumed constant for the sake of model simplification [16]. Therefore, as indicated by Eq. (S14), it is important to note that the competition coefficient *a_ij_* is proportional to the ratio of consumption rates *C* between consumer *j* and consumer *i*. Formally, we can represent it as

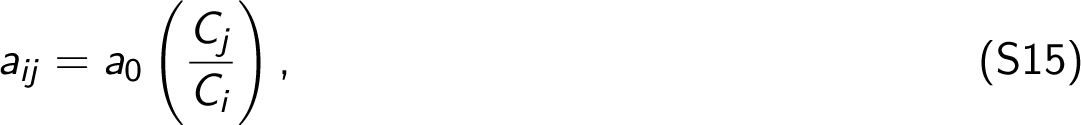

where *a*_0_ is a normalization constant. Following this, we know the intraspecific competition coefficient

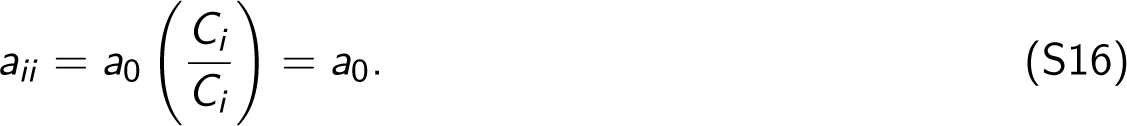

without loss of generality, we can set *a*_0_ = 1. Consequently, Eqs. (S12) become the LV competition equations:

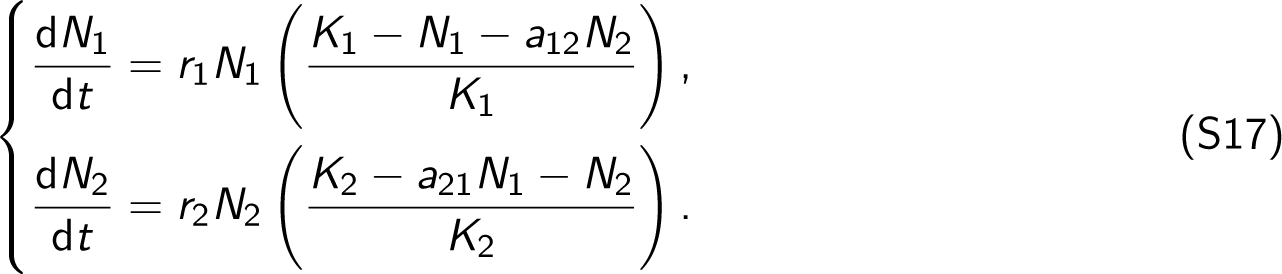

Considering the system *S* = *{*1, *· · ·*, *|S|}*, we can write the general form of the LV competition equations as

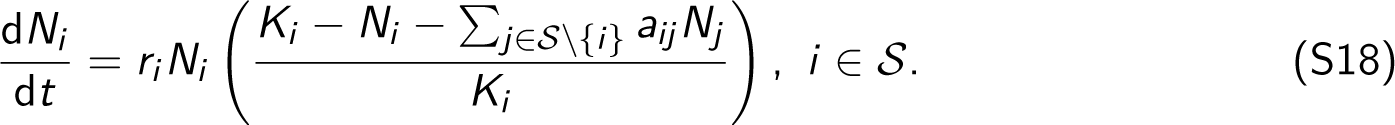

In the previous discussion, our focus has been on the scenario of two consumers and two resources. Next, we will proceed to simplify the scenario by considering only one consumer (also predator, either consumer *j* or consumer *i*) and one resource (also prey). Specifically, if the resource is self-replicating, then the resource equation with logistic growth is

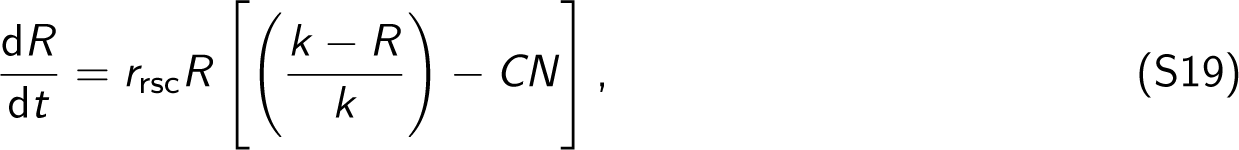

where *R* is the density of the resource, *r*_rsc_ is its intrinsic rate of increase, *k* is the carrying capacity of the resource, *C* is the rate at which the resource is consumed by the consumer, and *N* is the population density of the consumer.

Following the LV predator-prey equations, we can write the consumer equation as

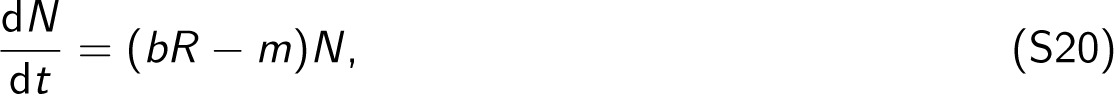

where *b* is the predation rate and *m* is the death rate of the consumer.

Assuming the resource dynamics are very fast compared to the consumer dynamics, similar to the idea in Eqs. (S6), we can solve the equilibrium value of R first:

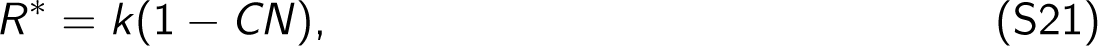

and then substitute *R^∗^* in place of *R* in the consumer equation Eq. (S20). We obtain

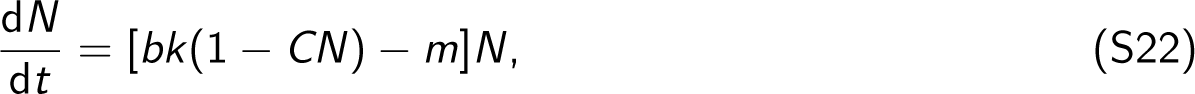

which can also be written as

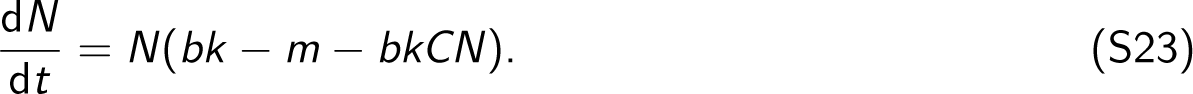

Multiplying and dividing Eq. (S23) by (*bk − m*) gives us

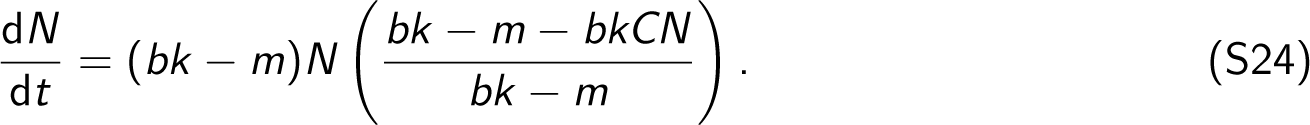

Next, dividing both the numerator and the denominator of the fraction on the right side of the Eq. (S24) by *bkC*, we have

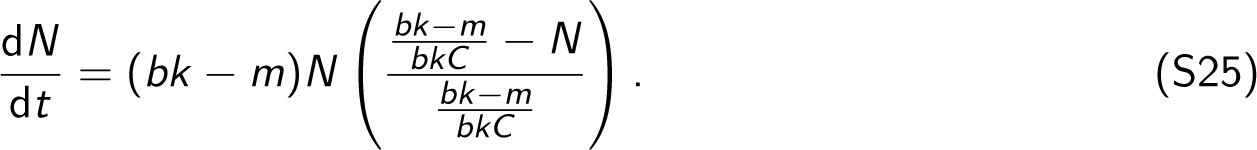

If we denote

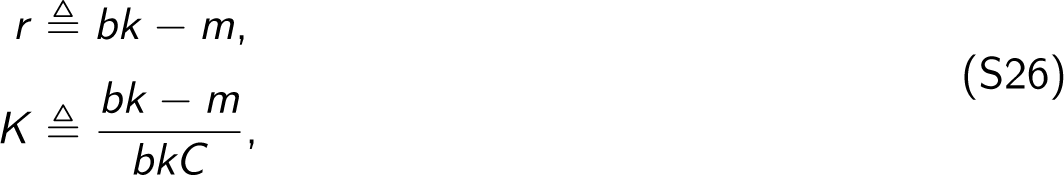

then Eq. (S25) can be written as the classic population model with logistic growth:

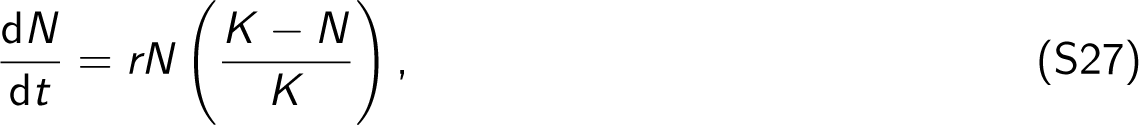

where *r* is the intrinsic rate of natural increase of the consumer in isolation and *K* is the carrying capacity of the consumer.

Importantly, Eqs. (S26) indicate that the consumption rate *C* increases monotonically with the intrinsic growth rate *r*. Formally, by substituting the expression for *bk* as (*r* + *m*) from the first equation into the second equation, we obtain

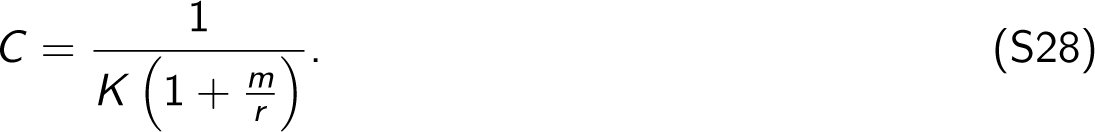

Fig. S19 shows the plots of Eq. (S28) with different death rates *m*, indicating that the consumption rate *C* exhibits a monotonically increasing behavior with respect to the intrinsic growth rate *r*. Also, the presence of an upper limit 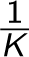 on *C* aligns with our empirical expectations, as it reflects the fact that the consumption rate cannot infinitely increase. Despite the relationship between *C* and *r* being nonlinear, it can be effectively approximated using a linear function, especially when *m* is large or comparable with the range of *r*. Note that the relationship between carrying capacity *K* and intrinsic growth rate *r* appears to be complex and context-dependent [17]. Therefore, instead of establishing relationships between *r* and *K*, our primary focus lies on *r* due to the relevant datasets available for microbial, plant, and metazoan systems. Nevertheless, if a negative relationship exists between *r* and *K*, such as 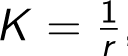, then *C* continues to increase monotonically with *r*. In this instance, Eq. (S28) transforms to 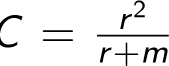, indicating a higher order of scaling in the relationship between *C* and *r*. This intensified relationship can potentially improve the performance of our theoretical framework. Exploring this additional complexity could be an interesting direction for future research.

### S4 Trade-offs induced by increasing growth rates (r)

Up to now, we have been discussing how to infer interactions *a_ij_* in the *K* -formalism of LV equations by linking them to the Consumer-Resource systems. Following the *r* -formalism of LV equations, it is evident that both interspecific and intraspecific interactions (*α_ij_*, *α_ii_*, respectively) tend to increase with growth rates *r*. Specifically, comparing the LV competition equation Eq. (S18) in *K* -formalism with the generalized LV model below in *r* -formalism

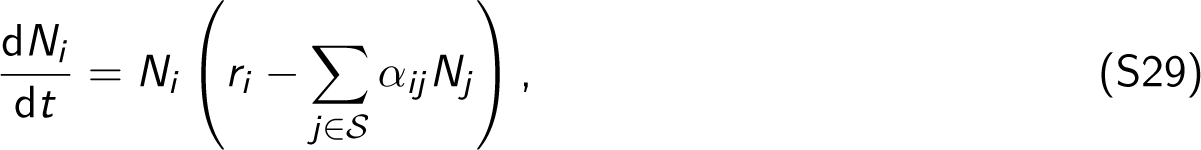

it becomes clear that

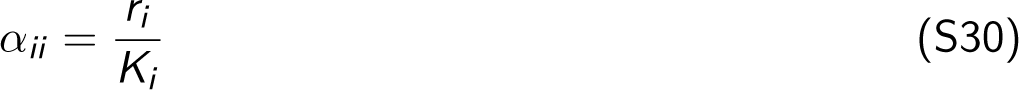

and

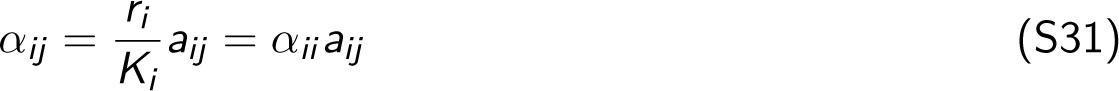

This relationship shows that the intraspecific interaction *α_ii_* is directly proportional to *r_i_*. In terms of the interspecific interaction *α_ij_*, we first deduce the expression of *a_ij_* explicitly integrating Eq. (S15) and Eq. (S28)

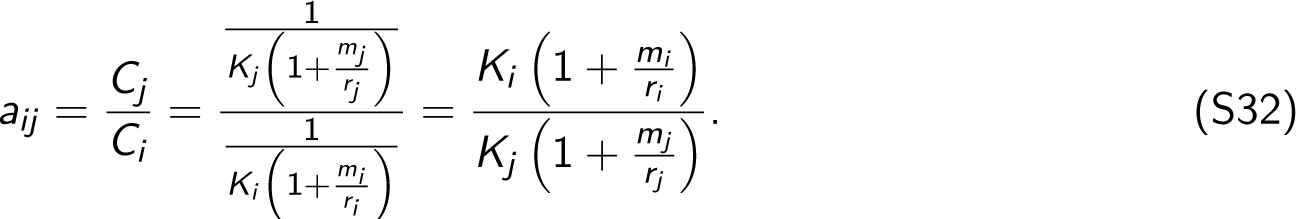

Then, following Eqs. (S30) and (S31), we have

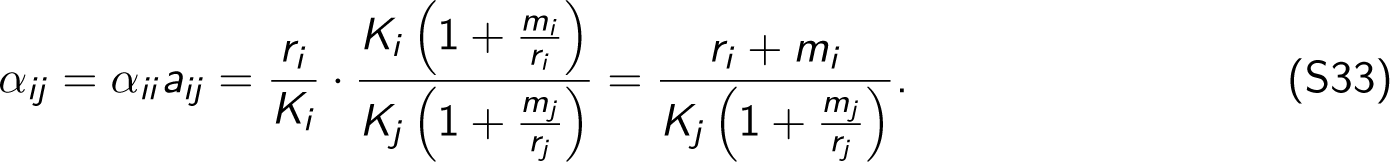

Therefore, the interspecific interaction *α_ij_* increases with growth rates *r_i_* and *r_j_* .

In summary, increases in growth rate translate into a trade-off where they carry increases in interspecific competition (*α_ij_*), but also they carry increases in density dependence (intraspecific competition *α_ii_*).

### S5 Inferring consumption rates (C) using maximum growth rates (rmax)

Next, based on Eq. (S28), the consumption rate *C* can be inferred to be approximately proportional to the intrinsic growth rate *r* (*C ∝ r*), which can be further approximated by the maximum growth rate (*r ≈* rmax). Formally, this relationship can be expressed as

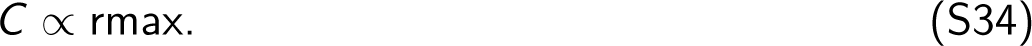

Here, we choose rmax as a parameter of interest because of its extensive investigation in empirical studies, particularly regarding its metabolic scaling laws. This choice allows for a systematic estimation of consumption rates *C* across a wide range of ecological systems, including bacteria, plants, and metazoans, leveraging the wealth of experimental datasets available.

Eventually, by combining Eq. (S15) with Eq. (S34), we can derive an estimation of competitive interaction strength *a_ij_* using the ratio of maximum growth rate (rmax) between consumer (taxon) *j* and consumer *i* (Fig. 1B):

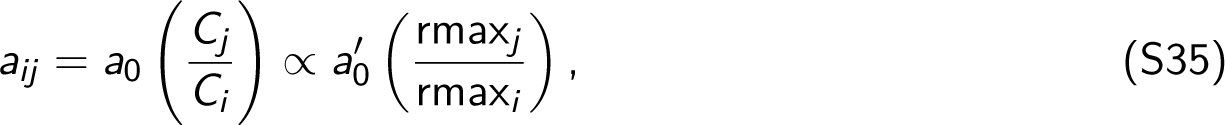

where *C_i_* and *C_j_* are the consumption rates of taxon *i* and taxon *j*, respectively; *a*_0_ and *a*_0_*^′^* are normalization constants. Based on this estimation, we can obtain the pairwise interaction matrix A of the Lotka-Volterra model (Eq. (S1)), which is the only parameter needed to implement the feasibility analysis. Phenomenologically, these competitive effects can represent either resource partition or modification [1–3, 18].

It is important to note that in our theoretical framework, we standardize parameters like carrying capacity (mentioned above) and dispersal ability (next section) to be equal across all taxa, allowing differentiation solely in their rmax. This level of control offers a coarse-grained explanation of the observed empirical patterns (see Figs. 3–4). Indeed, adjustments to these other parameters could further enhance the accuracy of the theoretical path of least resistance, enabling it to better fit the observed order of appearance in empirical systems compared to our current Fig. 3. This could be an interesting avenue for future research.

### S6 Quantification of the transition matrix (P) in the Markov chain

In the Markov chain of each developmental process, either sequentially or simultaneously, the transition matrix P plays a critical role as it summarizes the likelihood of transitioning from one ecological community to another. Mathematically, the transition matrix of a developmental process is a square matrix, where the rows and columns represent the ecological communities characterized by different taxon compositions. The matrix size is equal to the number of all possible ecological communities in the developmental process. Each element in the matrix represents the probability of transitioning from the community corresponding to the row to the community corresponding to the column.

Here, we consider bottom-up developmental processes that allow for loops wherein a community may transition back to itself. Specifically, communities comprising *n* taxa can either transition to communities comprising (*n* + 1) taxa with an additional taxon available in the taxon pool, or they can loop back to themselves and remain unchanged. This coarse-grained model suffices to provide an overall expectation of taxon appearance orders in ecological development. Note that considering cycles (e.g., disassemblies caused by environmental perturbations), which allow a larger community to collapse into a smaller one, does not affect our theoretical conclusions regarding the path of least resistance (see Sec. S8).

The likelihood of community transitions within an ecological system is determined by the feasibility of all possible communities within it. Below is a detailed example using the three-taxon system *S* illustrated in Fig. 1, which comprises taxa tx1, tx2, and tx3 (i.e., *S* = *{*123*}*). We will systematically explore the quantification of transition matrices in the Markov chains that describe sequential development from initial communities with one or two taxa, as well as simultaneous development from the community containing all three taxa. Then the three transition matrices are presented. Using these matrices, we can calculate the cumulative probability over *x* trials for developing an ecological system.

Initially, all developmental processes start from an empty set ∅. Note that, as mentioned before, we focus on the order in which taxa appear during development, rather than the specific composition of resulting communities. Thus, this empty set provides a standardized baseline, allowing for meaningful comparisons of appearance orders across various types of development, regardless of whether a system starts with no taxa or undergoes a reset due to disturbances. Subsequently, the system may either remain in the empty set or transition into communities containing varying numbers of taxa. On the one hand, the probability of remaining empty is the complement of the average feasibility of all possible initial communities. If the initial system contains one taxon (Fig. 1D), the probability that the system remains empty is 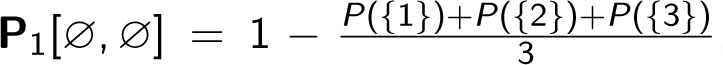, where *P*(*{*1*}*), *P*(*{*2*}*), and *P*(*{*3*}*) represent the feasibility of the communities with tx1, tx2, and tx3, respectively (see Eq. (S3)). Notably, the likelihood of any single taxon being feasible in unknown environments is 50% (i.e., *P*(*{*1*}*) = *P*(*{*2*}*) = *P*(*{*3*}*) = 0.5), similar to flipping a fair coin. This equal feasibility implies that all taxa have the same dispersal ability and are not restricted to appear in a specific order because of this. Instead, their order of appearance is driven by their growth rates and the current subset of present taxa (their interactions). If the initial system contains two taxa, then the probability of remaining empty is 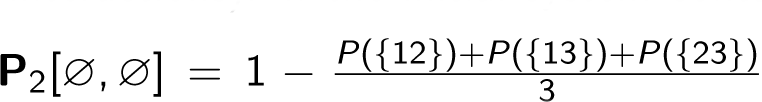, where *P*(*{*12*}*), *P*(*{*13*}*), and *P*(*{*23*}*) represent the feasibility of the communities with pairs tx1 and tx2, tx1 and tx3, and tx2 and tx3, respectively. If all three taxa appear initially and simultaneously, the probability of remaining in the empty set is P_3_[∅, ∅] = 1 *− P*(*{*123*}*), where *P*(*{*123*}*) is the feasibility of all three taxa coexisting together in system *S* (see Fig. 1E). On the other hand, the probabilities of transitioning to different initial communities are proportional to their respective feasibility. Importantly, by the definition of a Markov chain, these probabilities collectively sum to 1 *−* P[∅, ∅]. For example, the transition probabilities from the empty set ∅ to the three one-taxon initial communities satisfy P_1_[∅, *{*1*}*] : P_1_[∅, *{*2*}*] : P_1_[∅, *{*3*}*] = *P*(*{*1*}*) : *P*(*{*2*}*) : *P*(*{*3*}*) and P_1_[∅, *{*1*}*] + P_1_[∅, *{*2*}*] + P_1_[∅, *{*3*}*] = 1 *−* P_1_[∅, ∅]. For initial communities containing two taxa, the transition probabilities from empty satisfy P_2_[∅, *{*12*}*] : P_2_[∅, *{*13*}*] : P_2_[∅, *{*23*}*] = *P*(*{*12*}*) : *P*(*{*13*}*) : *P*(*{*23*}*) and P_2_[∅, *{*12*}*] + P_2_[∅, *{*13*}*] + P_2_[∅, *{*23*}*] = 1 *−* P_2_[∅, ∅].

For intermediate communities, a community may either remain unchanged or transition into larger communities by incorporating an additional taxon. The probability of remaining in the current community equals the feasibility of that community; for example, P_1_[*{*1*}*, *{*1*}*] = *P*(*{*1*}*). Alternatively, the probabilities of transitioning into larger communities are proportional to their respective feasibility. For instance, a community with tx1 may transition to a larger community by including an additional taxon in the pool, either tx2 or tx3. The probabilities for these transitions are proportional to their feasibility, namely, P_1_[*{*1*}*, *{*12*}*] : P_1_[*{*1*}*, *{*13*}*] = *P*(*{*12*}*) : *P*(*{*13*}*). These transition probabilities are located in the row of *{*1*}* and the columns of *{*12*}* and *{*13*}* within the transition matrix P_1_. Similarly, for the community comprising tx1 and tx2, the probability of maintaining the current composition is equivalent to the feasibility of the community, *P*(*{*12*}*). Alternatively, this community may transition into a larger community that includes tx3, the only remaining candidate in the taxon pool, with a complementary probability of (1 *−P*(*{*12*}*)). These transition probabilities are then cataloged in the row of *{*12*}* and the columns of *{*12*}* and *{*123*}* within the transition matrix P_1_. In terms of intermediate communities, the transition probabilities for developmental processes that start with two or more taxa (e.g., P_2_) can be quantified in the same way as those described above for processes starting with a single taxon.

Lastly, the transition probability for the community consisting of tx1, tx2, and tx3 (all taxa in the pool) returning to itself is 1. This probability is saved in the row and column *{*123*}* of the transition matrices P_1_, P_2_, and P_3_. This indicates that the full community *{*123*}* is an absorbing state. Once all taxa are assembled, the community does not undergo further changes, serving as an endpoint in the sequence of community changes. Note that this absorbing state is the sufficient (but not necessary) condition for development. If there is interest in exploring beyond the endpoint community to understand how additional taxa might integrate, it is also possible to delay reaching an endpoint by expanding the absorbing state. For example, this can be done by increasing the number of taxa in the initial pool and analyzing the dynamics of the intermediate communities that assimilate these additional taxa of interest.

The Markov chain allows both sequential and simultaneous developments to be quantified using the same set of rules, establishing a basis for their comparability. Notably, the simultaneous development, due to its simple structure, can be analytically shown to be equivalent to a binomial process (refer to Sec. S7 for more details). For instance, the transition matrix P_1_ for developing a three-taxon system sequentially from one-taxon communities can be formally represented as

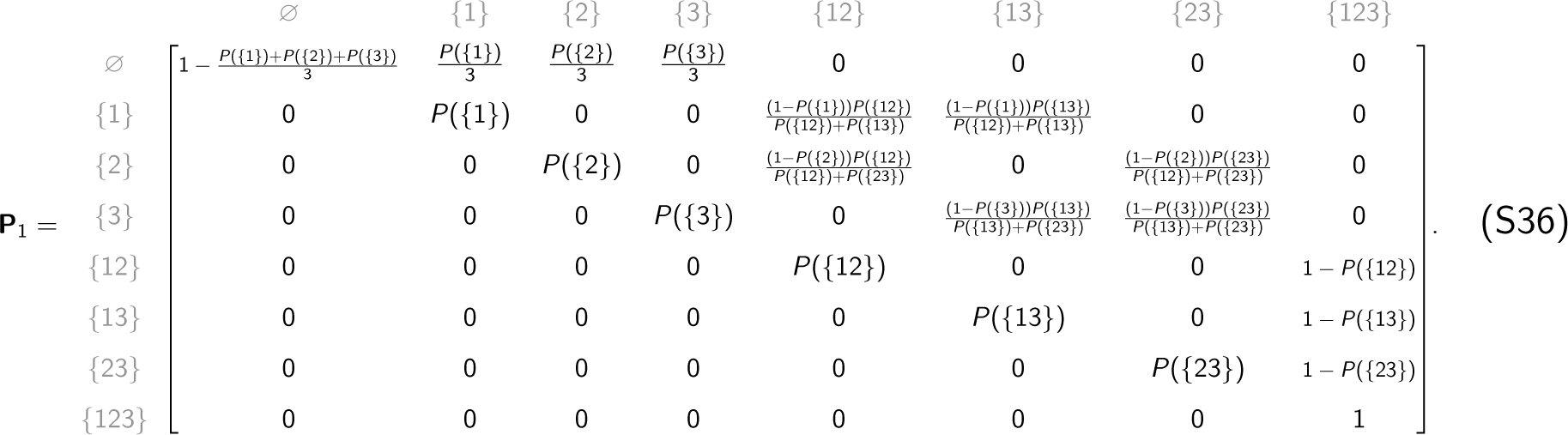

Similarly, the transition matrix P_2_ for developing a three-taxon system sequentially from two-taxon communities can be represented as

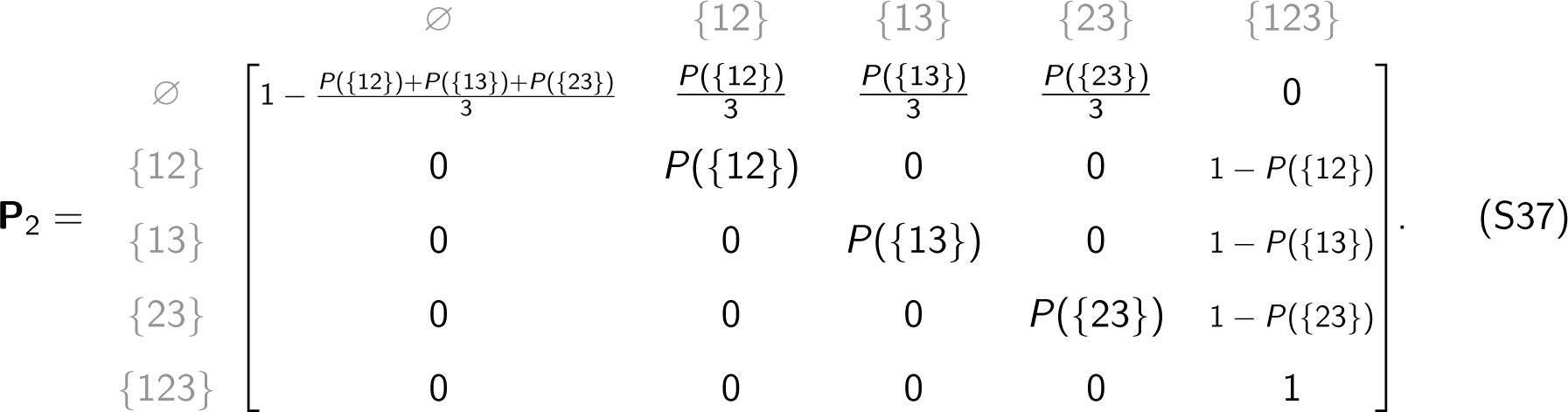

In simultaneous development, the transition matrix P_3_ for developing a three-taxon system can be represented as

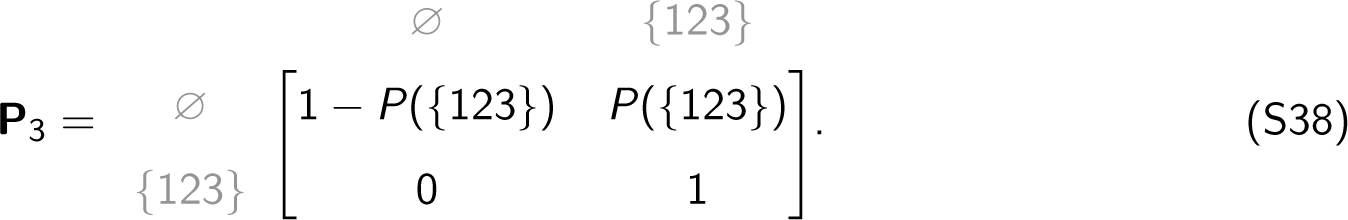

A key property of the Markov chain is that raising the transition matrix P to the power of *x* yields the probabilities of transitions between ecological communities with *x* steps (or trials). Based on this property, we can calculate the cumulative probability P*^x^* [∅, *{S}*] of developing an *n*-taxon ecological system *S* over *x* attempts through sequential processes with varying numbers of initial taxa or the simultaneous process (see Eqs. (S36)-(S38); Fig. 1F). Moreover, since *{S}* is an absorbing state, the cumulative probability P*^x^* [∅, *{S}*] across all developmental processes would always converge to probability 1 with sufficiently large numbers of trials. However, the rate of convergence can vary depending on the sizes of the initial communities. Therefore, it is possible to identify the process with the highest likelihood of successfully developing the entire system within a limited number of trials (see results in Figs. 2A, S2A, and S4A). In short, the larger the initial communities, the slower the convergence. Thus, as detailed in the following section, convergence to probability 1 occurs more slowly in simultaneous development compared to sequential development.

### S7 Understanding simultaneous development as a binomial process

Let P be the transition matrix of a Markov chain describing the simultaneous development of an *n*-taxon ecological system *S*. Analogous to the transition matrix of a three-taxon system in Eq. (S38), the general form for an *n*-taxon system is

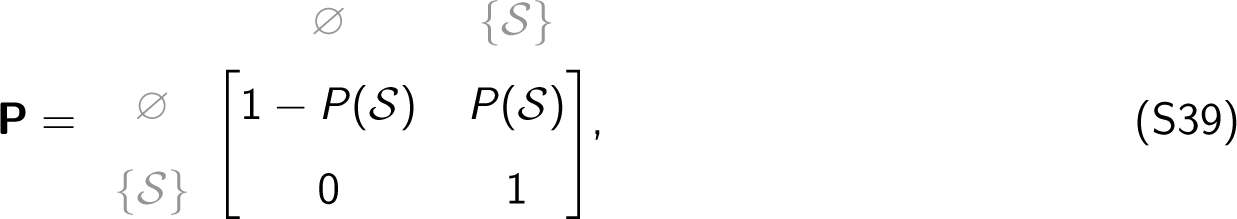

where P[∅, *{S}*] = *P*(*S*) denotes the probability of transitioning from the empty set ∅ to the ecological system *S* (see Fig. 1E). This transition probability is quantified as the feasibility of the *n*-taxon system, indicating the likelihood of achieving simultaneous coexistence (see Eq. (S3)).

We propose that the *x* -step transition matrix P*^x^* is given by the following formula for all integers

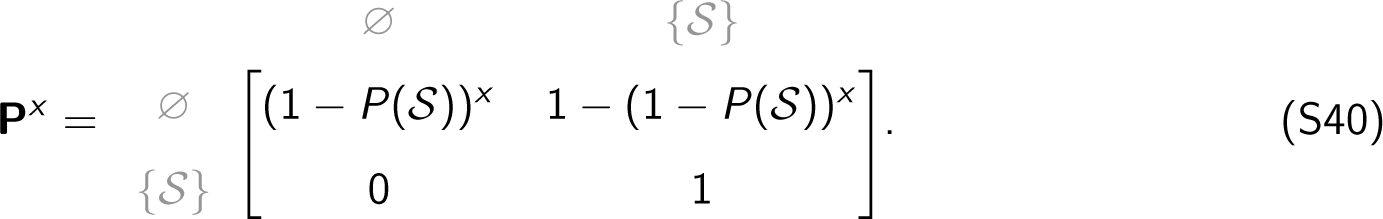

This formula will be proven by mathematical induction, starting with the base case when *x* = 1. The one-step transition matrix P^1^ is simply

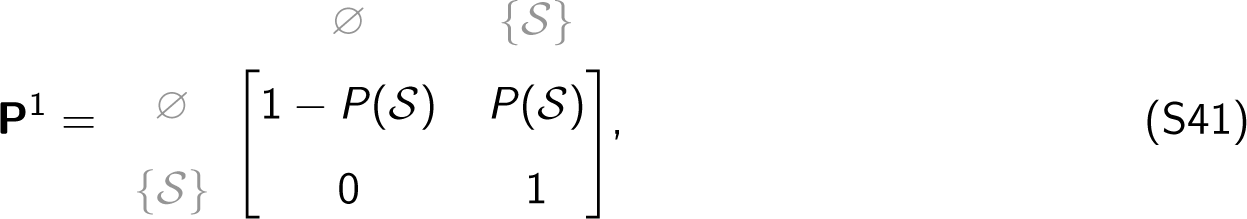

which matches the given matrix in Eq. (S39). Thus, the base case holds.

Assuming inductively that the formula Eq. (S40) holds for some arbitrary integer *x ≥* 1, we proceed to show that this formula also holds for (*x* + 1). The (*x* + 1)-step transition matrix P*^x^*^+1^ can be computed by multiplying P*^x^* by P. Thus,

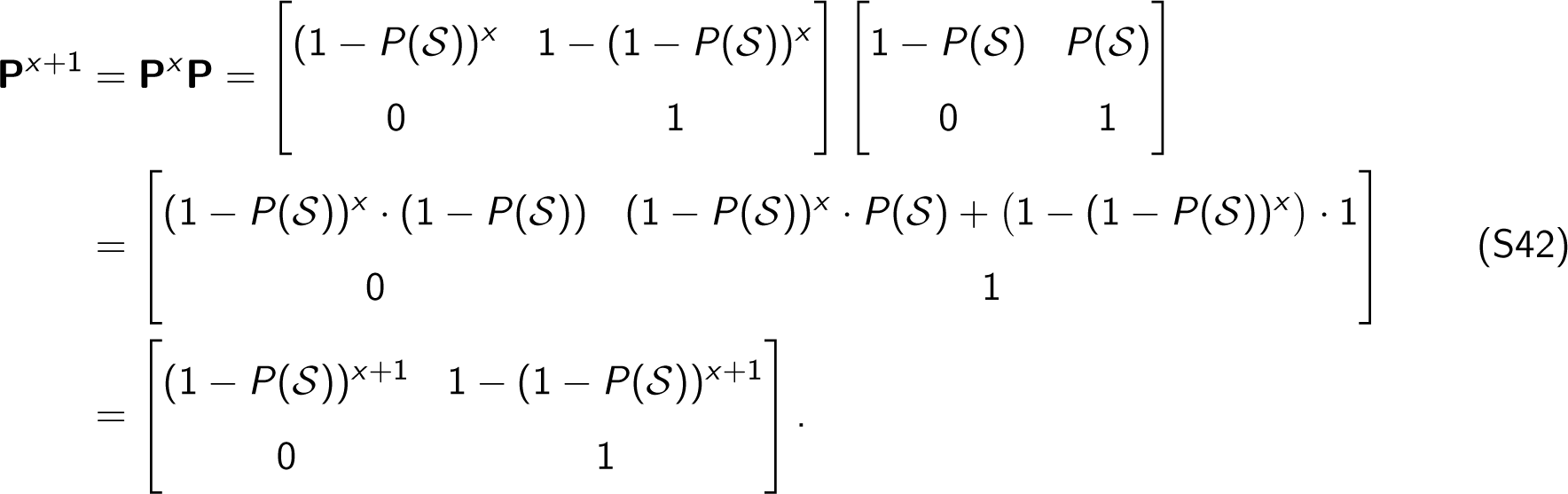

Clearly, the formula Eq. (S40) holds for (*x* + 1). By induction, this confirms that for all integers *x ≥* 1, the *x* -step transition matrix P*^x^* is given by Eq. (S40).

Taking a closer look at Eq. (S40), it is interesting to notice that the simultaneous process can be characterized not only as a Markov chain (Eq. (S39)) but also as a binomial process, similar to flipping a coin. Specifically, consider the analogy of flipping a biased coin, where

- Heads (success) corresponds to transitioning to the system *S*. This occurs with a probability of *P*(*S*).
- Tails (failure) corresponds to remaining in the empty set ∅. This occurs with a probability of 1 *− P*(*S*).

If the coin is flipped *x* times (representing *x* steps in the Markov chain), then the probability of flipping at least one head (transitioning to system *S* at least once) during these *x* attempts is 1 *−* (1 *−P*(*S*))*^x^*. Note that (1 *−P*(*S*))*^x^* is the probability of the complement event, which is getting all tails (never transitioning to system *S* in any of the *x* attempts). As *x* increases, this complement probability approaches zero because the effect of repeatedly multiplying a number less than one diminishes its value towards zero. Thus, 1 *−* (1 *− P*(*S*))*^x^* converges to 1, indicating that with a sufficiently large number of attempts, transitioning to system *S* through simultaneous development becomes almost certain. However, since the probability of all taxa coexisting *P*(*S*) is very low (more taxa lead to harder coexistence and thus a lower probability), the convergence of simultaneous development is significantly slower compared to sequential development (Figs. 2A, S2A, and S4A).

Therefore, the binomial process offers an intuitive way to understand the inherent stochastic processes that influence the simultaneous development of an ecological system. It quantifies the likelihood that the system successfully reaches simultaneous coexistence at least once within a given number of attempts.

### S8 Identification of the path of least resistance

We define development as the self-organized process of moving from an empty set to a set of *n* taxa through a collection of different taxon communities (interactions) whose union forms the set of *n* taxa (i.e., the appearance of a group of taxa). In the main text, we focus our analysis on the special case of directional assembly (once a taxon is successfully introduced it cannot go extinct). Effectively, this analysis aims to explain the probability that a group of *n* species can be found together at a given point in time. While this analysis is a simplification of the developmental process undergone by natural systems (i.e., no extinctions); under our assumptions, this analysis generates the same results as if additionally considering disassembly processes (see Sec. S9). That is, under assembly only, the question is which one path is the system most likely to take. Under assembly and disassembly, the question would be which is the most visited path by the system.

Formally, a directional assembly (one-by-one bottom-up developmental process) of an *n*-taxon ecological system *S* is a sequence of community changes ∅, *C*_1_, *C*_2_, *· · ·*, *C_n−_*_1_, *S* consisting of 0, 1, 2, *· · ·*, *n −* 1, *n* taxa, respectively (for every *i* such that 1 *≤ i ≤ n −* 1, *C_i_ ⊂ S*). In a Markov chain, the probability of observing this sequence can be calculated by multiplying the initial probability with the conditional probabilities of each transition:

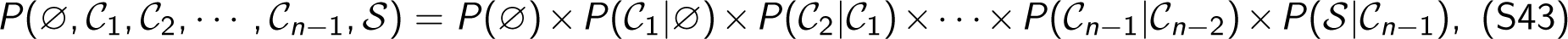

where *P*(∅) = 1 is the probability of developing from an empty set. This standardized baseline, with equal probabilities for all paths, allows for meaningful comparisons. In the context of full environmental uncertainty (no a priori information), the probability *P*(*C*_1_*|*∅) of having a single taxon is analogous to flipping a fair coin, with a constant 50% chance of success. Ecologically, this constant probability suggests that all taxa have the same likelihood of appearing. Also, this probability can be adjusted in light of prior environmental information [3]. Moreover, it is important to note that we do not assume any relationship between a taxon’s dispersal probability and its maximum growth rate. However, should there be a positive relationship, it would result in an even higher probability *P*(∅, *C*_1_, *C*_2_, *· · ·*, *C_n−_*_1_, *S*) for the path of least resistance, thus enhancing the performance of our theoretical framework and the most likely appearance order from fast-to slow-growing taxa. Incorporating this additional complexity offers a compelling direction for future research. Additionally, *P*(*C_m_|C_m−_*_1_) is the transition probability from community *C_m−_*_1_ to community *C_m_* (*m ≤ n*, *C*_0_ = ∅, *C_n_* = *S*), which is the element located at the intersection of row (*m −* 1) and column (*m*) in the transition matrix P. This equation holds because the Markov chain has the memoryless property that a future system depends only on the current system, not on any preceding systems in the sequence. The effects of memory in ecological developmental processes could be an interesting avenue for future research.

To identify the path of least resistance, the objective is to find the sequence of ecological systems that maximizes the probability *P*(∅, *C*_1_, *C*_2_, *· · ·*, *C_n−_*_1_, *S*). This can be conveniently achieved by transforming the multiplication operation in Eq. (S43) into a summation operation. Specifically, this transformation can be achieved by applying the negative logarithm function (*−* log) to both sides of the equation. Any base greater than 1 is valid, including common bases such as 10, *e*, and 2. That is,

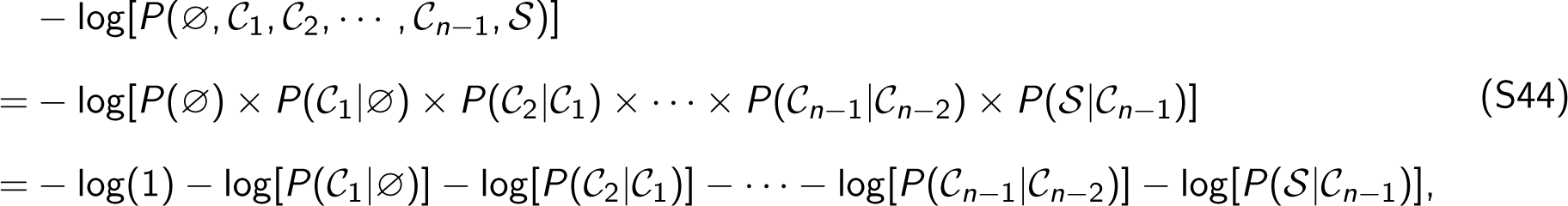

where *P*(∅) = 1 is a constant. The R codes accompanying this paper use the negative natural logarithm function (*−* ln) with a base of *e*.

The objective now shifts to minimizing *−* log[*P*(∅, *C*_1_, *C*_2_, *· · ·*, *C_n−_*_1_, *S*)], which is essentially minimizing 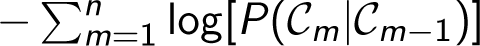, where *C*_0_ = ∅ and *C_n_* = *S*. In graphical terms, our objective is to find the shortest path in the deterministic tree-like directed graph characterized by the adjacency matrix *−* log P. In this graph, vertices represent ecological communities of different taxon compositions. Besides, each edge carries a weight *−* log[*P*(*C_m_|C_m−_*_1_)] that represents the distance from its connected vertex *C_m−_*_1_ to the other connected vertex *C_m_*. In the context of ecological development, the shortest path is from a vertex representing the empty set ∅ to a vertex representing the most complex community *S*. According to graph theory, Dijkstra’s algorithm provides an efficient solution to this shortest path problem [19]. The R codes provided use the function “shortest paths” from the “igraph” package for these computations. The path with the shortest distance among all paths is defined as the path of least resistance for developing an ecological system (e.g., pink water pipes in Fig. 1E and pink arrows in Fig. S3).

All computational tasks were executed using the High-Performance Computing resources available through the MIT SuperCloud and the Lincoln Laboratory Supercomputing Center [20]. The R codes are accessible on GitHub at: https://github.com/MITEcology/Deng_Assembly.

### S9 Theoretical predictions are not influenced by disassembly

As mentioned in the main text, our analysis focuses on the special case of directional assembly, wherein once a taxon is successfully introduced, extinction is precluded. Though this assumption simplifies the natural developmental process, it effectively serves our purpose of determining the expected order of appearance of taxa. It is important to note our theoretical predictions remain consistent even when disassembly processes are considered additionally. With assembly only, the question is which path the system is most likely to follow. Conversely, with both assembly and disassembly, the question is which path is the most frequently visited path by the system.

In this section, we will start by developing ecological intuitions to understand why disassembly does not affect our theoretical predictions. Following this, we will incorporate randomly generated disassembly and repeat the analysis presented in Fig. 2. This allows us to systematically and quantitatively assess the influence of disassembly.

In the special case of directional assembly only (see Fig. 1D), sequential development from smaller initial communities can diverge into multiple paths. This makes it more efficient to reach the largest community compared to those with a greater number of initial taxa. Even with disassembly present (Fig. S3), these finely bifurcated paths from smaller initial communities can still expedite the progression towards the largest community. Therefore, it can be anticipated that the cumulative probability of sequential development from a single taxon reaches its saturation faster than others with more initial taxa, including simultaneous development (i.e., assembling all taxa in the pool at once). Additionally, a developmental process progresses in a single direction in the case of assembly only, resulting in a saturation probability of 1 (*y* -axis in Fig. 2A). This indicates that once taxa are introduced, the system inevitably advances towards the formation of the largest community without any reversals. However, when disassembly processes are incorporated, a new possibility arises. Disassembly introduces the potential for species loss and regression to less developed communities, disrupting the progression towards the largest community. Hence, the cumulative probability of reaching the largest community is expected to fall below 1. Essentially, the inclusion of disassembly introduces additional opportunities for deviations from the most likely developmental trajectory, leading to a lower cumulative probability of development.

Interestingly, despite the potential impact of disassembly, the most likely developmental order (the path of least resistance) remains unaffected. Let us take the transition matrix P in Eq. (S36) as an example, which describes the sequential development for a three-taxon system sequentially from one-taxon communities.

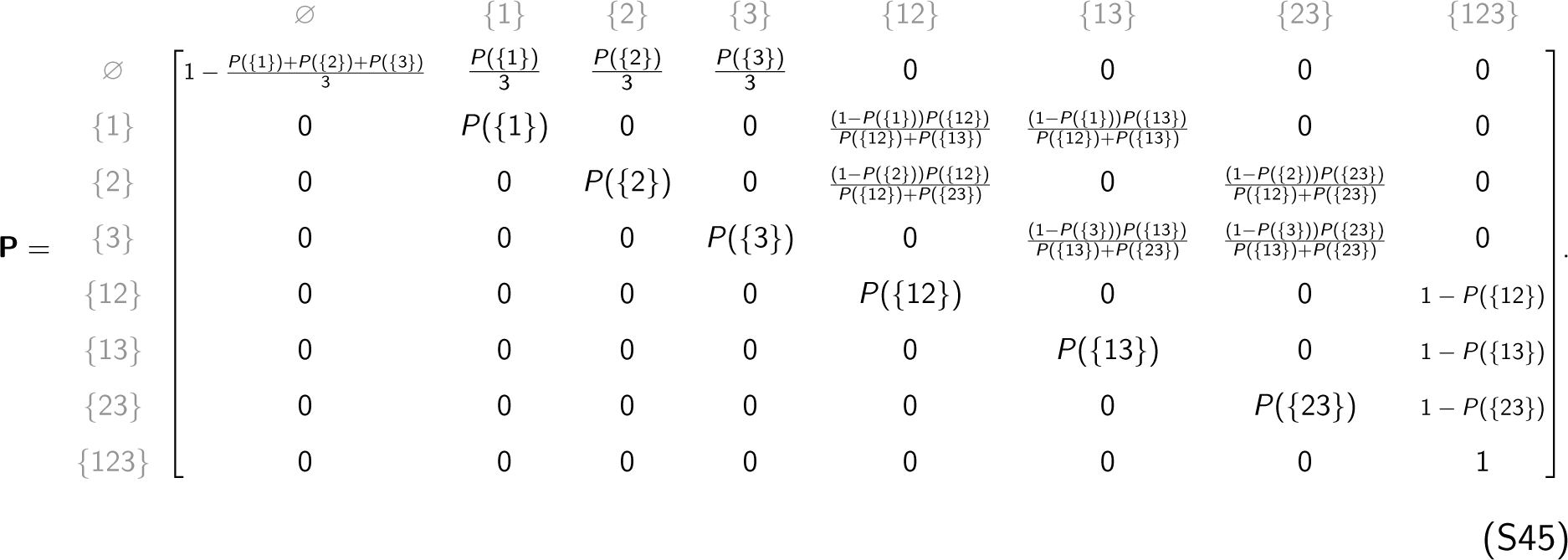

By definition, each element in the transition matrix is the probability of transitioning from the community in the row to the community in the column. The upper triangular portion represents the assembly process of an ecological system, and the lower triangular portion represents the disassembly process. Recall that the path of least resistance is defined as the path that is the most likely path for the system to take. Let us assume that ∅ *→ {*1*} → {*12*} → {*123*}* is the path of least resistance considering the special case of directional assembly only. That is, the probability *P*(∅, *C*_1_, *C*_2_, *{*123*}*) of observing the sequence of community changes from ∅ to *{*123*}*, i.e., P[∅, *C*_1_] *×* P[*C*_1_, *C*_2_] *×* P[*C*_2_, *{*123*}*], can be maximized when *C*_1_ = *{*1*}* and *C*_2_ = *{*12*}* (see Eq. (S43)). Note that P[∅, *{*1*}*], P[*{*1*}*, *{*12*}*], and P[*{*12*}*, *{*123*}*] are all located in the upper triangular portion of the transition matrix (see Eq. (S45)) and therefore influenced solely by the assembly process, not the disassembly process. Ecologically speaking, though assembly and disassembly processes are often intertwined in natural systems, the most likely order in which taxa appear for biodiversity development remains independent of disassembly processes driven by uncertain environmental perturbations. This can be intuitively understood by recognizing that the path of least resistance, or the shortest path, is by nature the most straightforward route, which tends to avoid any return cycles to a less developed community as a result of disassembly (see Fig. S3).

With these intuitions in mind, we aim to systematically verify whether our theoretical predictions in Fig. 2 remain valid when additionally considering disassembly. To do this, we randomly generated disassembly probabilities for each community. These random probabilities account for the fact that different communities may disassemble with varying probabilities in response to uncertain environmental perturbations. Note that incorporating disassembly mathematically means filling the lower triangular portion of a transition matrix (e.g., Eq. (S45)) with non-zero values. Specifically, these disassembly probabilities were drawn from a uniform distribution ranging from a minimum of 0 to a maximum of 0.1. The results remain robust to other maximum values. To ensure the validity of the studied developmental process, we enforced that the assembly probability for each transition must be greater than the reverse disassembly probability, thus allowing overall biodiversity to increase. In cases where this condition was not met, we iteratively generated another disassembly probability from the same distribution mentioned above.

Using the transition matrix incorporating both assembly and disassembly processes, we conducted the same analysis as presented in Fig. 2A. We computed the cumulative probability for development from the empty set ∅ through varying initial communities to the full community *S* (refer to Sec. S6 for more details). The results are presented in Fig. S4A, indicating that sequential developments starting from smaller initial communities tend to reach saturation more quickly. Notably, in all developments, the cumulative probability of reaching the largest community falls below 1 (grey dashed line), which is in line with our expectations above. Even with disassembly present, sequential development remains more likely to succeed than simultaneous development. These results are consistent with our theoretical prediction in Fig. 2A.

Although the most likely developmental paths begin with communities comprising a single taxon, various sequences with different orders of appearance are possible. For example, both sequences ∅ *→ {*1*} → {*12*}* and ∅ *→ {*2*} → {*12*}* start from a single taxon but introduce the two taxa in opposite orders. It is therefore important to identify the single most likely path. To this end, we used Dijkstra’s algorithm to find the most likely path, also referred to as the path of least resistance (see Sec. S8 for more details). With this path established, we conducted the same analysis as presented in Fig. 2B. The results, presented in Fig. S4B, reveal that when disassembly is considered, the expected correlations between the most likely appearance order of a taxon and its rmax are generally negative (predominantly *y <* 0; Spearman’s rank correlation). Also, as the standard deviation of rmax among taxa (*x* -axis) increases, these negative correlations tend to be stronger (approaching *y* = *−*1). These results are consistent with our theoretical prediction in Fig. 2B.

In summary, with disassembly processes additionally considered, our theoretical predictions shown in Fig. 2 remain consistent. Furthermore, the most likely developmental order along the path of least resistance is not altered by disassembly. Thus, the theoretical most likely order for each empirical system presented in Figs. 3-4 remains the same.

### S10 Proxy for maximum growth rate (rmax)

In this paper, we investigate six published empirical datasets of varying ecological systems, including infant gut microbes [21], marine bacteria [22], trees [23], birds [24], and herbivores [25, 26]. To estimate the maximum growth rate (rmax) for each system, we employ metabolic scaling laws, which have been providing valuable insights into the fundamental processes that govern biological structure and function.

Specifically, Reference [5] presents the scaling relationships between rmax and body mass across prokaryotes (including bacteria and archaea) and metazoans (including birds and herbivores). Interestingly, the authors discover a shift in the scaling pattern of rmax from positive in prokaryotes (with a power of 0.73 *≈* 3*/*4) to negative in metazoans (with a power of *−*0.23 *≈ −*1*/*4). Therefore, following the scaling relationship rmax *∝* (body mass)*^−^*^1^*^/^*^4^, we use the body mass information reported in the AVONET database [27] and References [25, 26] to estimate the rmax for birds and herbivores, respectively (Figs. 3D–3F).

Similarly, for plants, Reference [6] demonstrates an inverse scaling relationship between rmax and body mass, characterized by a power of *−*1*/*4. Additionally, Reference [28] provides an allometric scaling relating tree mass (M) to diameter at breast height (dbh) via the formula *M* = 78 *× dbh*^2.47^. Consequently, to approximate the rmax for individual tree species, we use diameter at breast height data from the BIEN database [29–33] (Fig. 3C).

However, due to the considerable variation in microbial body mass under environmental uncertainties and limited reporting, we use the ribosomal RNA operons (rrn) copy number, rather than the body mass, as a proxy for rmax in microbial systems. Reference [7] presents the positive scaling relationship between the rrn copy number and the rmax of bacteria and archaea, i.e., rmax *∝* (rrn copy number)^5^*^/^*^4^. Also, the authors supply a comprehensive database of rrn copy numbers called rrnDB [34], from which we obtain the average rrn copy numbers for the six phyla of infant gut microbes and the five orders of marine bacteria (Figs. 3A–3B).

It is important to note that the choice of scaling exponent does not affect the qualitative results of our framework. Let us explain this with the following example. Considering that the body mass distribution *M*_1_, *M*_2_, *· · ·*, *M_n_* of *n* taxa in an ecological system follows a log-normal distribution with a log mean of *µ* and a log standard deviation of *σ*, then it is clear that log(*M*_1_), log(*M*_2_), *· · ·*, log(*M_n_*) are normally distributed with a mean of *µ* and a standard deviation of *σ*. Assuming that a scaling relationship exists between body mass and rmax, this can be represented as rmax*_i_* = *M_i_^α^*, where *α ∈* (*−∞*, *∞*) is a constant scaling exponent. This scaling relationship can also be expressed logarithmically as log(rmax*_i_*) = log(*M_i_^α^*) = *α ·* log(*M_i_*) for each taxon *i* within the range 1, *· · ·*, *n*. Thus, log(rmax_1_), log(rmax_2_), *· · ·*, log(rmax*_n_*) are also normally distributed with a mean of *αµ* and a standard deviation of *|α|σ*, where *| · |* denotes absolute value. By definition, this means that rmax_1_, rmax_2_, *· · ·*, rmax*_n_* of the *n* taxa follow a log-normal distribution with a log mean of *αµ* and a log standard deviation of *|α|σ*. Therefore, the scaling exponent *α*, regardless of its sign (due to *| · |*) or magnitude, does not alter the log-normal nature of the distribution of maximum growth rates (rmax).

### S11 Details of empirical data

The first empirical system of infant gut microbes (presented in Figs. 3A and 4A; [21]) was drawn from a case study that tracked the gut microbial composition of a healthy male infant over 2.5 years, beginning on day 3. From a time series of more than 60 fecal samples, this dataset represents primary succession (*de novo* system development) and consists of six main phyla of bacteria and archaea (observed order of appearance): Firmicutes (#1), Proteobacteria (#2), Actinobacteria (#3), Bacteroidetes (#4), Verrucomicrobia (#5), and Euryarchaeota (#6). This observed order of appearance is shown in Figure 6A and Table S6 of Reference [21].

The second empirical system of marine bacteria (presented in Figs. 3B and 4B; [22]) was derived from a laboratory microcosm study. The authors immersed chemically defined, nutrient-rich microparticles in coastal seawater over a span of 144 hours. Then they recorded a collection of bacterial strains isolated from different phases of colonization and generated 16S rRNA gene sequences. This dataset also represents primary succession and consists of five main orders of bacteria (observed order of appearance): Alteromonadales (#1), Vibrionales (#2), Flavobacteriales (#3), Oceanospirillales (#4), and Rhodobacterales (#5). This observed order of appearance is shown in Figure 3A of Reference [22].

The third empirical system of trees (presented in Figs. 3C and 4C; [23]) was from a 500-acre woods in Princeton, NJ, owned by the Institute for Advanced Study. The forest was disturbed by a blight in 1920 and hurricanes in 1938 and 1944. Given the slow pace of forest succession, which makes direct observation impractical, the author inferred the successional patterns, including the order of taxon appearance, using the ages of all trees on various plots of land with unknown histories. The author has also confirmed these inferred successional patterns with further observations. This dataset represents secondary succession (local reassembly of an established species pool following an environmental change) and consists of 7 species (order of appearance): Betula populifolia (#1), Liquidambar styraciflua (#2), Populus grandidentata (#3), Nyssa sylvatica (#4), Acer rubrum (#5), Liriodendron tulipifera (#6), and Fagus grandifolia (#7). This order of appearance is shown in Figure SUCCESSIONAL CHANGES IN PRODUCTIVITY on Page 95 of Reference [23].

The fourth empirical system of birds (presented in Figs. 3D and 4D; [24]) was from abandoned agricultural fields in the Piedmont region of Georgia, United States. Between 1947 and 1951, the authors counted and identified birds across ten areas at various stages of natural vegetative succession, ranging from one-year abandoned fields to young mature forests aged over 150 years. This dataset represents secondary succession and consists of 14 main families (observed order of appearance): Parulidae (#1), Passerellidae (#1), Cardinalidae (#1), Vireonidae (#2), Polioptilidae (#3), Paridae (#3), Troglodytidae (#3), Trochilidae (#4), Sittidae (#4), Tyrannidae (#4), Picidae (#4), Turdidae (#4), Corvidae (#4), and Cuculidae (#5). This observed order of appearance is shown in Table 1 of Reference [24]. The ruby-throated hummingbird (*Archilochus colubris*), the only representative of Trochilidae in the dataset, is a known specialist that feeds largely on nectar from certain types of flower and sustains one of the highest mass-specific metabolic rates of any vertebrate [35]. Because the unique foraging behavior and outlying physiology of this species relative to others in the dataset has the potential to bias our inferences (e.g., by skewing detection probability in the original data depending on the timing and location of surveys in relation to flowering phenology), we excluded it from the main text analysis as an outlier. However, we obtained qualitatively similar results when including the hummingbird (see Fig. S5).

The fifth empirical system of herbivores (presented in Figs. 3E and 4E; [26]) was from Gorongosa National Park, in central Mozambique, southeast Africa. The data reflect a 28-year time series of aerial count data on ungulates and elephants from 16 genera (*Tragelaphus* being the only genus with *>*1 species) from 1994 to 2022 [25]. Surveys were conducted both before (6 surveys of 9 genera from 1968–1972) and after (14 surveys of all 16 genera from 1994–2022) the Mozambican Civil War (1977–1992), which caused *>*90% declines in all large-herbivore taxa [25]. Densities were determined by dividing the number of animals counted by the maximum area surveyed in each year (1968–2018 data are from Table S3 of Reference [25]; 2020 data are from Reference [36], and 2022 data are from Reference [37]). The 14 postwar surveys analyzed here were conducted at varying intervals (1–3 years) using different methodologies (3 fixed-wing, 11 by helicopter) at different scales (106–2195 km^2^). The earlier surveys included the three fixed-wing counts (1994, 1997, 2004) and were conducted at smaller scales (average 472 km^2^ from 1994–2007 vs. 1705 km^2^ from 2010–2022). To account for sampling artifacts arising from these discrepancies, we calculated moving averages of densities in windows of 3 successive surveys (e.g., the value calculated for 2020 is the average of the 2018, 2020, and 2022 counts). This yielded estimates for 12 postwar intervals. We note that smaller species are harder to detect in aerial surveys, especially in fast-flying fixed-wing aircraft, but this bias is conservative with respect to our prediction (i.e., smaller species are less likely to be detected, particularly in early years). Because low observed densities do not necessarily indicate a viable population, we set three density thresholds (25%, 50%, and 75%), each relative to the maximum density recorded for the genus across all pre- and postwar surveys; a genus is classified as appeared when its density exceeds the threshold for the first time. These thresholds signify different assumptions about when a taxon can be considered to have successfully (re)established inside the park, as opposed to being vagrant individuals or nascent populations vulnerable to stochastic extinction. Taxa that are increasing in density but have not reached a given threshold were assigned the last order of appearance. For example, in Fig. S6A, wildebeest (*Connochaetes*), buffalo (*Syncerus*), and zebra (*Equus*) were all present in 2022 but had not yet met the 25% threshold. The implicit assumption that these taxa *will* arrive is justified by their rapidly increasing densities: in fact, wildebeest exceeded 25% of maximum prewar density in 2022, but this is not captured in our moving-window approach, while buffalo (which are threefold larger with a correspondingly lower rmax) reached 11% and have been increasing by an average of 12% yr*^−^*^1^ since 2016. The only uncertainty is zebra (*>*2% of prewar maximum in 2022 and still at risk of failure), but excluding zebra does not alter the result. Notably, the reassembly of large herbivores in Gorongosa is an overwhelmingly natural recovery process (the only genera appreciably supplemented by translocations are *Connochaetes*, *Syncerus*, and *Equus*, which have not yet arrived under any of our thresholds) and has been relatively robust to external perturbations [37, 38]. In the main text (Figs. 3E and 4E), we report results using a conservative 25% threshold. Fig. S6 shows the results under all three thresholds.

The sixth empirical system of herbivores (presented in Figs. 3F and 4F; [26]) was from Gorongosa National Park. We analyzed encounter frequencies with individuals of different large mammalian herbivores (ungulates and elephants, 17–3825 kg) while driving road transects in the south of the park during the progression from the early to late dry season (June–August) of 2016. These data reflect secondary succession in response to seasonal change within a *∼*250 km^2^ focal area and consist of 11 genera (observed order of appearance): *Aepyceros* (#1), *Kobus* (#1), *Phacochoerus* (#2), *Alcelaphus* (#3), *Ourebia* (#4), *Tragelaphus* (#5), *Redunca* (#6), *Hippotragus* (#7), *Loxodonta* (#8), *Connochaetes* (#9), and *Syncerus* (#10). Again, only one genus comprises *>*1 species (*Tragelaphus* with 3 closely related, ecologically similar species [39]). The observed order of appearance is from the metadata deposited in Dryad Digital Repository accompanying Reference [26]. The seasonal results are consistent with the multi-decadal analysis (Figs. 3E, 4E, and Fig. S6) and provide support for our theory over extraordinary organismal and spatiotemporal scales.

For the microbial systems, we extracted rrn copy numbers from the rrnDB database [34], from which we estimated the rmax of infant gut microbes and marine bacteria. In these two systems, the rmax of each taxonomic rank, either a phylum or an order, was calculated as the average rmax estimated by the average rrn copy number across all documented species in the rrnDB database within the respective taxonomic level. For the forest system, we used the diameter at breast height data from the BIEN database [29–33] to estimate the rmax of each species. For the animal systems, we used body mass data from the AVONET database for bird species [27] and Reference [26] for herbivore species to estimate the rmax (Fig. 1B). The rmax for higher taxonomic ranks, either a family or a genus, was calculated based on the average body mass across all species in the respective group. We then assessed the correlation between rmax and observed appearance order using Spearman’s rank correlation coefficient (*ρ*). Similarly, we used Spearman’s *ρ* to evaluate the correlation between the observed path and the theoretically predicted path of least resistance and compared this with the distribution of *ρ* values obtained from 10^3^ randomly generated paths (see results in Fig. 3 and Figs. S5A, S6A–S6E).

To test our theoretical predictions about the influence of diversity in rmax on developmental order, we drew subsets of taxa from the entire empirical system to generate smaller assemblages with varying degrees of diversity in rmax. We then compared Spearman’s rank correlations between rmax and observed order of appearance across these subsets, segmenting diversity (standard deviations) of rmax into different groups (see results in Fig. 4 and Figs. S5B–S5D, S6B–S6F). For illustration, we delineated these subsets as 60% of the entire system size and the number of diversity groups as three. However, our results are robust to variation in these parameters (see Figs. S7–S18).

## S12 Supplementary figures

**Supplementary Figure S1:**
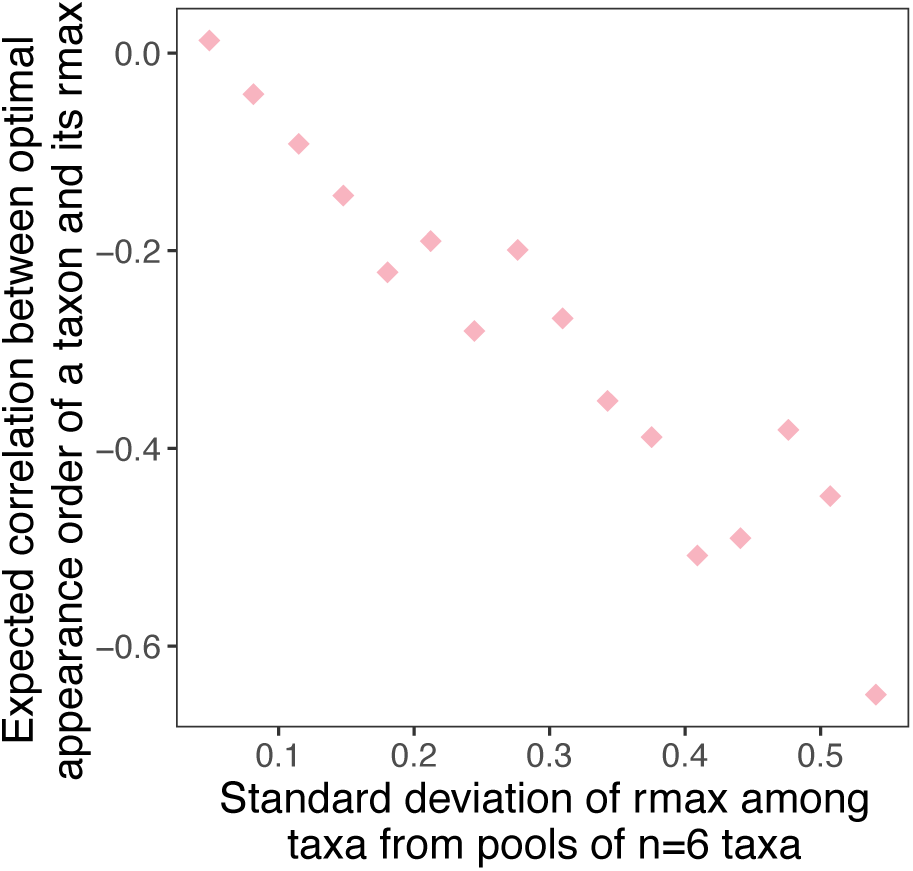
Reproducing Fig. 2B in the main text using a power of -1/4. Recall that the power in Fig. 2B is 3*/*4, i.e., rmax *∝* (body mass)^3^*^/^*^4^. Here we use rmax *∝* (body mass)*^−^*^1^*^/^*^4^. Similarly, the expected correlations between the most likely appearance order of a taxon and its rmax are generally negative (*y <* 0). Also, as the standard deviation of rmax among taxa increases, these negative correlations tend to be stronger (approaching *y* = *−*1). This observation confirms that the results remain consistent regardless of the value of the power in the scaling law (refer to Sec. S10 for a detailed explanation).

**Supplementary Figure S2:**
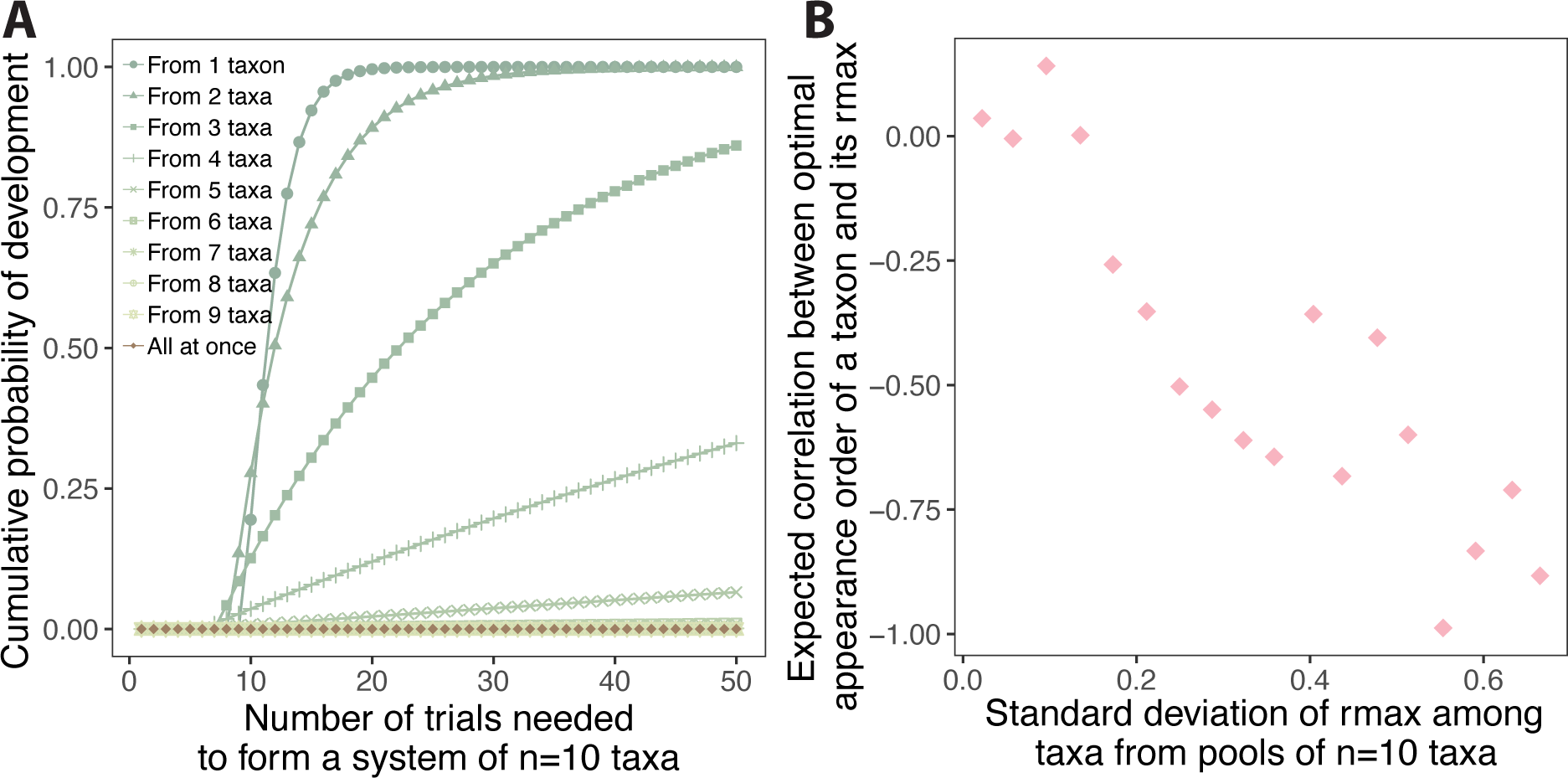
Reproducing Fig. 2 in the main text using systems of n = 10 taxa. (A) The sequential development starting from one taxon (darker green circles) reaches probability 1 more quickly compared to those starting with larger initial communities. Also, the cumulative probability of development through the simultaneous process (brown) is negligibly small, though it may reach probability 1 if the number of attempts is sufficiently large (Secs. S6 and S7). (B) The expected correlations between the most likely appearance order of a taxon and its rmax are generally negative (*y <* 0). As the standard deviation of rmax among taxa (*x* -axis) increases, these negative correlations tend to be stronger (approaching *y* = *−*1). This observation confirms that the results remain consistent regardless of system sizes. The power relationship here is rmax *∝* (body mass)^3^*^/^*^4^. The results are qualitatively consistent across different power relationships (see Figs. 2B and S1).

**Supplementary Figure S3:**
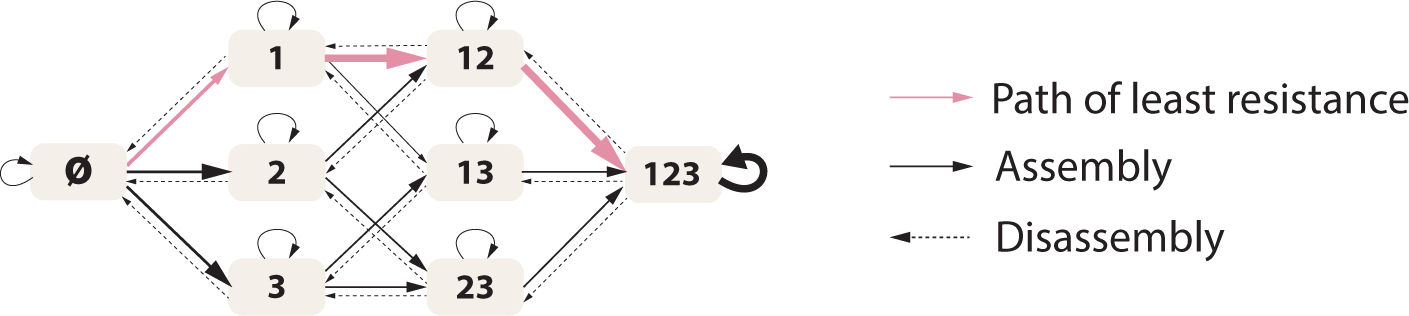
The path of least resistance in ecological development is not influenced by disassembly. The path of least resistance is exclusively determined by the assembly process in the development of an ecological system and remains unaffected by the presence or speed of disassembly, despite that these two processes are often intertwined in nature. This can be easily understood by recognizing that the path of least resistance, or the shortest path, is defined as the most straightforward route, which tends to avoid any return loops to a less developed community as a result of disassembly (see Sec. S9).

**Supplementary Figure S4:**
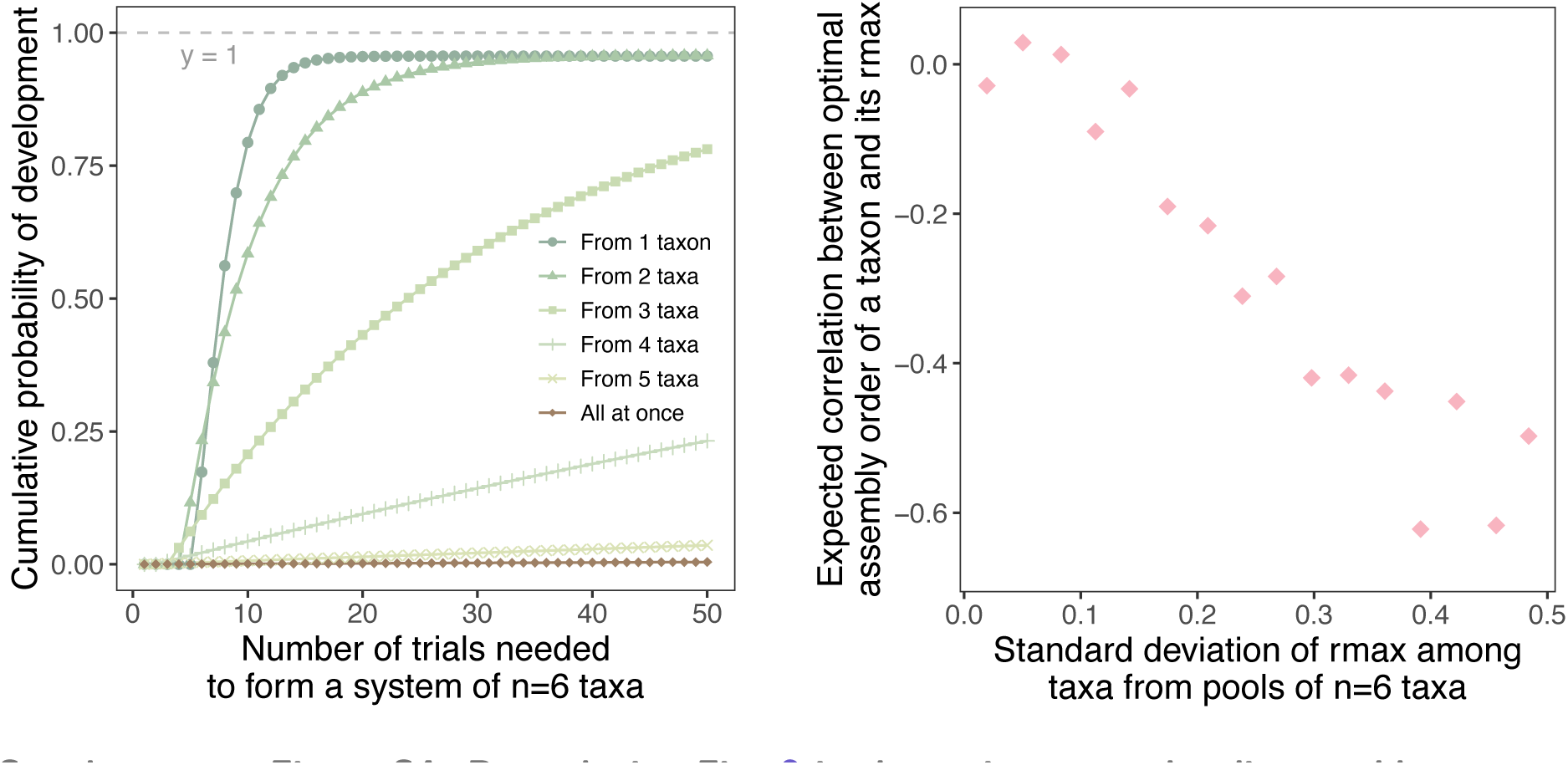
Reproducing Fig. 2 in the main text under disassembly. (A) The sequential development starting from one taxon (darker green circles) reaches its saturation probability more quickly compared to those starting with larger initial communities and simultaneous development. Note that under disassembly, the saturation probability is less than 1 (grey dashed line). This occurs because disassembly introduces the possibility of taxa being removed from the community, thus reducing the overall probability of reaching the largest community. (B) The expected correlations between the most likely appearance order of a taxon and its rmax are generally negative (*y <* 0). As the standard deviation of rmax among taxa (*x* -axis) increases, these negative correlations tend to be stronger (approaching *y* = *−*1). Both panels confirm that the results remain consistent when additionally considering disassembly. Please refer to Sec. S9 for more details.

**Supplementary Figure S5:**
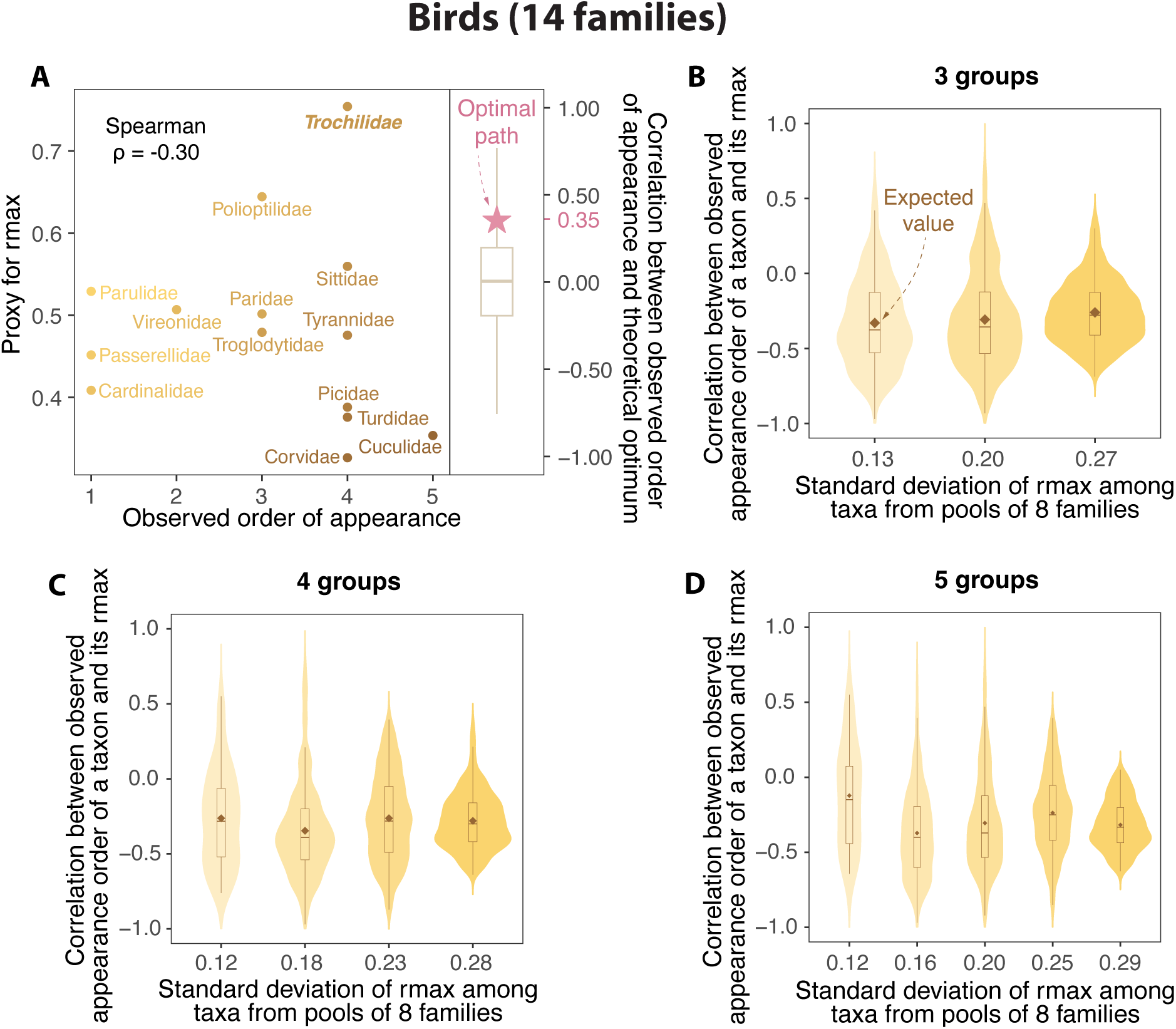
Reproducing Fig. 3D and Fig. 4D in the main text using the bird dataset that includes the Ruby-throated Hummingbird (Trochilidae: *Archilochus colubris*) (A) The Spearman’s rank correlation between the observed order of appearance (left; *x* -axis) and the proxy for rmax (*y* -axis) is *ρ* = *−*0.30. The Spearman’s rank correlation between the observed order of appearance and the theoretical order of the path of least resistance (right; pink star) is *ρ* = 0.35. Despite the unique feeding behavior (and associated potential sampling bias) and outlying mass-specific metabolic rate of hummingbirds, the analysis including this species is qualitatively consistent with that shown in Fig. 3D. (B)–(D) The correlations between the observed appearance order of a taxon and its rmax are negative (*y <* 0). However, an increase in the standard deviation of rmax among taxa (*x* -axis) does not consistently strengthen the negative correlations (*y* approaches *−*1). This reflects the influence of the high inferred rmax and late inferred appearance of hummingbirds. Here we choose the pool size as 60% of the entire system size. We cluster standard deviations of rmax into 3, 4, and 5 different groups to show their distribution. Please refer to Sec. S11 for more details about this dataset.

**Supplementary Figure S6:**
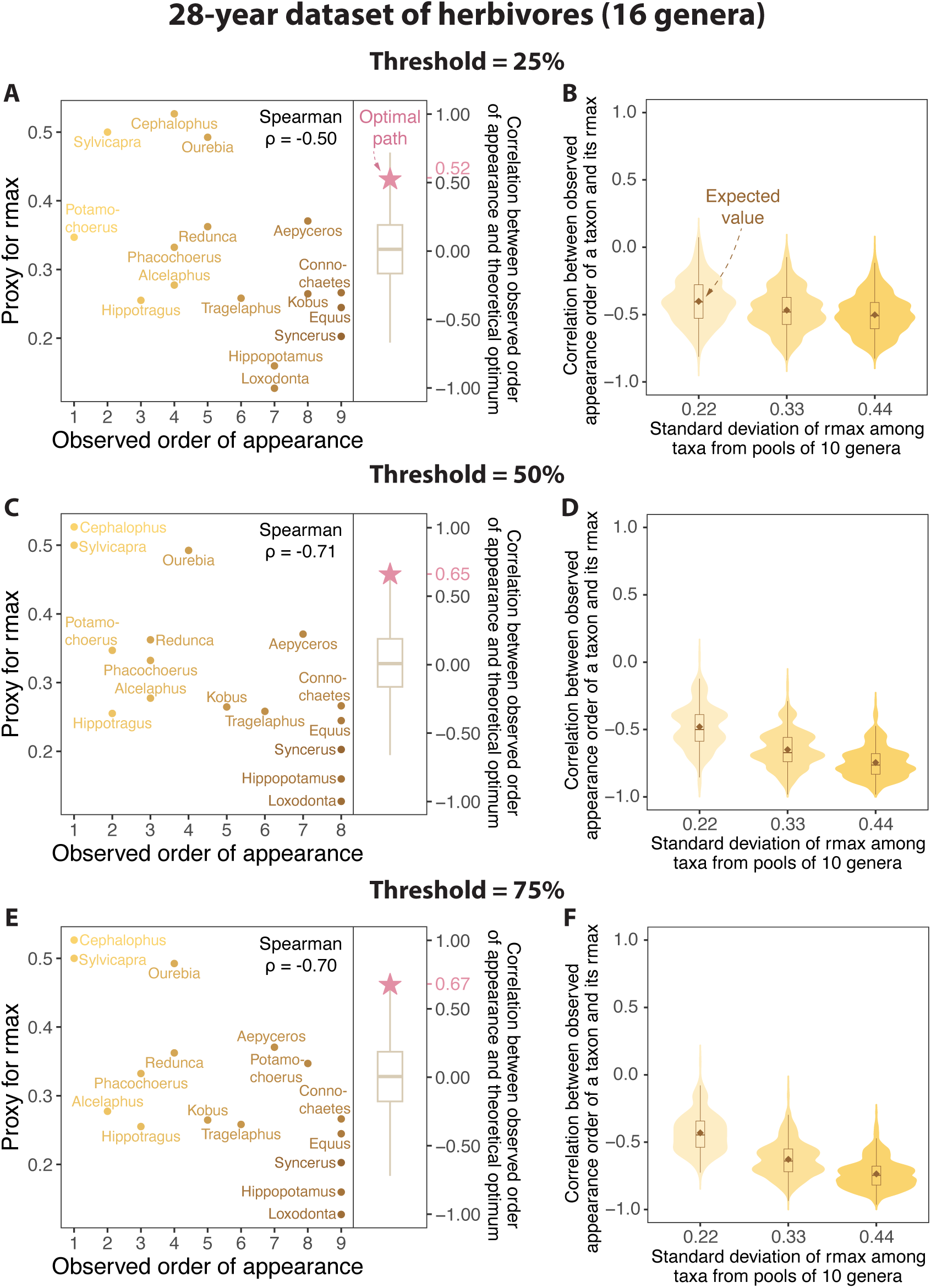
Postwar re-development of herbivores in Gorongosa National Park, based on aerial counts from 1994 to 2022. The observed orders of appearance of herbivore genera are based on density thresholds set at (A)–(B) 25% (presented in the main text), (C)–(D) 50%, and (E)–(F) 75% of the maximum density observed for each taxon either before or after the Mozambican Civil War. Taxa are classified as arrived when they first surpass a given density threshold. The results remain consistent regardless of threshold selection. Details on this dataset are in Sec. S11.

**Supplementary Figure S7:**
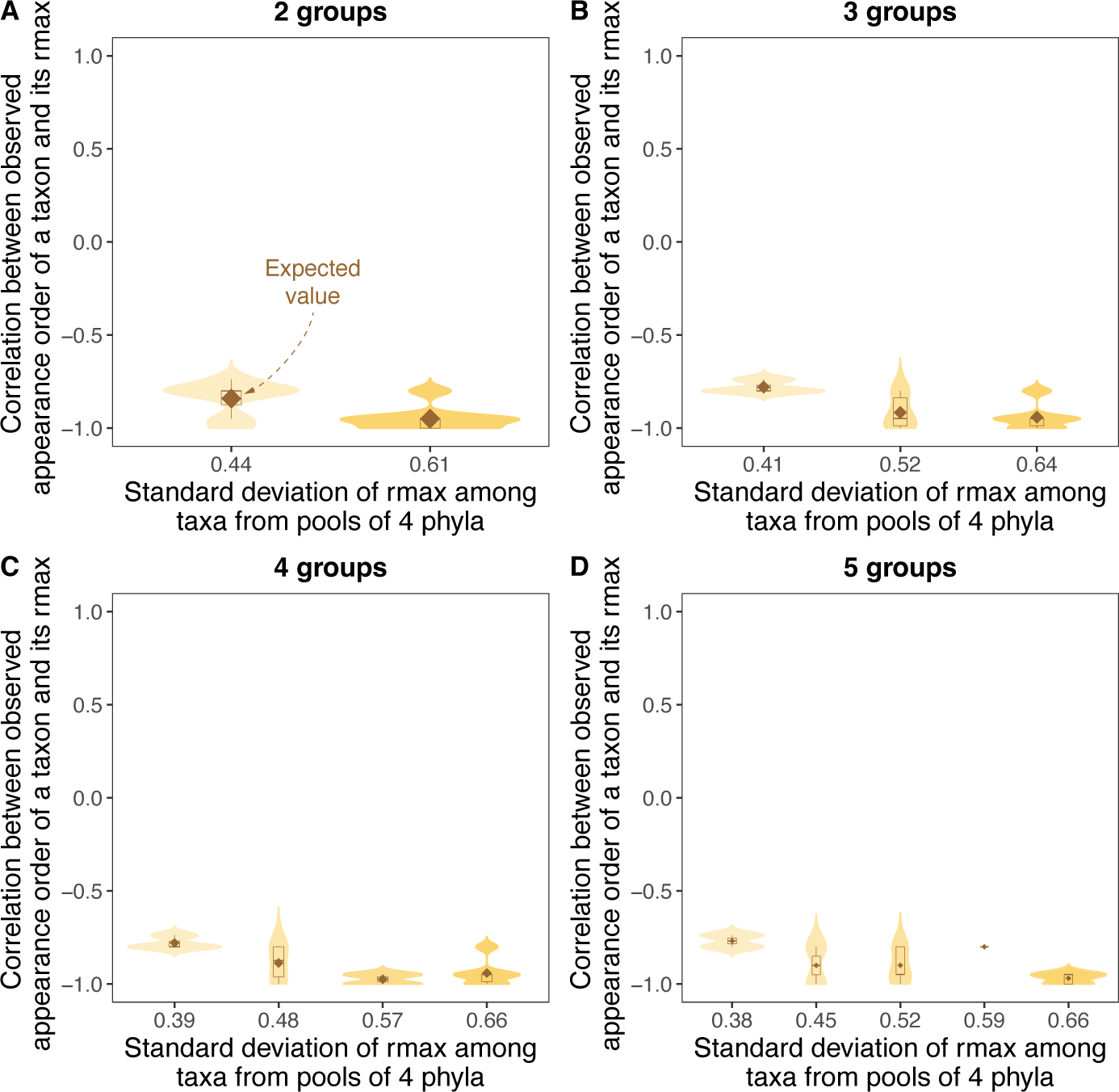
Infant gut microbes 4 phyla pools (subsets) clustered into different numbers of groups based on standard deviations of rmax. We cluster standard deviations of rmax into 2, 3, 4, and 5 different groups to show their distribution. The results remain consistent with the theoretical prediction in Fig. 2B regardless of group numbers. Details on this dataset are in Sec. S11.

**Supplementary Figure S8:**
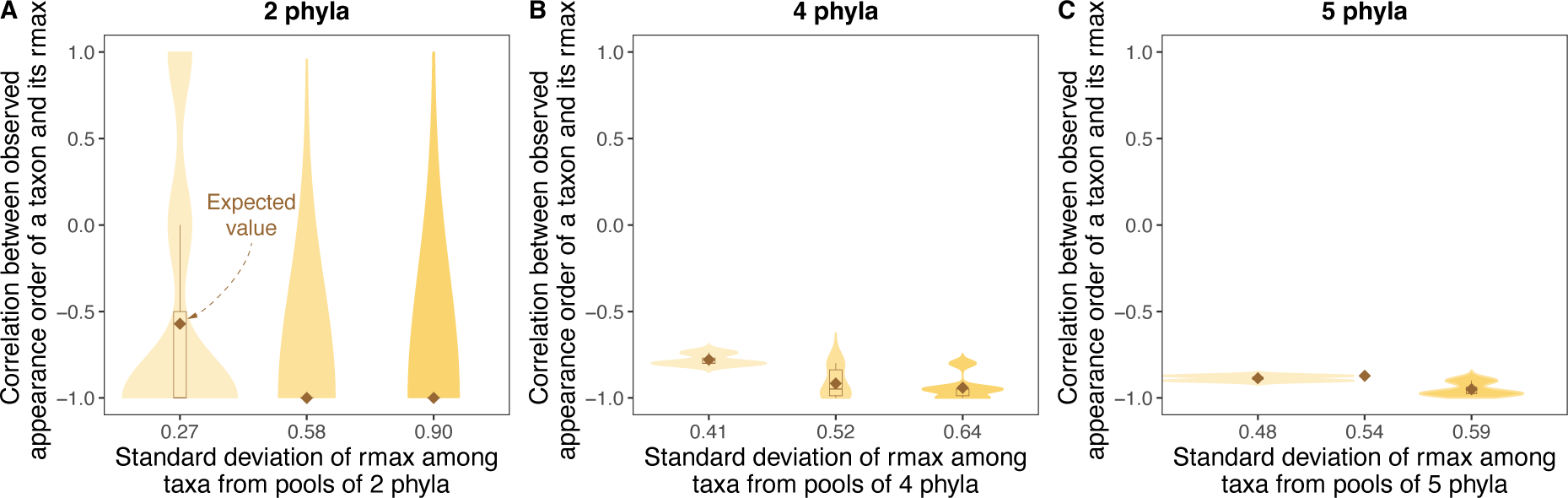
Infant gut microbes pools (subsets) with different numbers of phyla. We select pool sizes that are 40%, 60%, and 80% of the entire system size of 6 phyla. The results remain consistent with the theoretical prediction in Fig. 2B regardless of pool sizes. Details on this dataset are in Sec. S11.

**Supplementary Figure S9:**
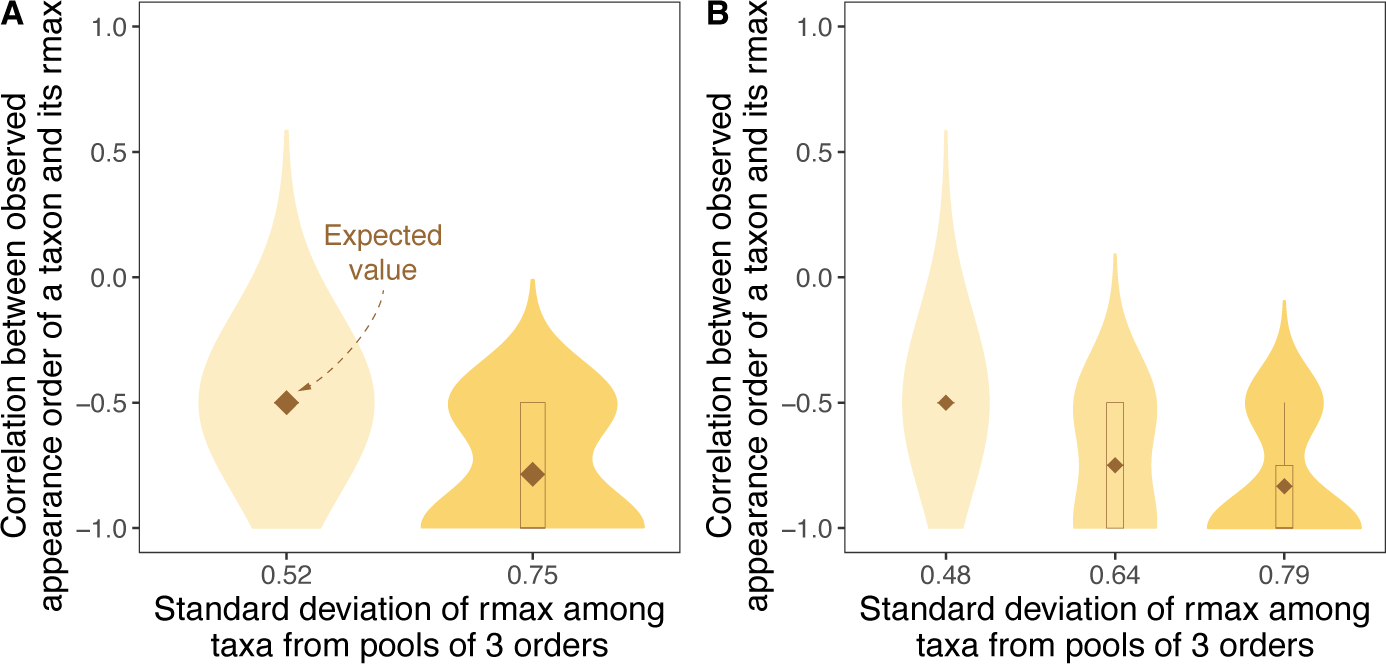
Marine bacteria 3 orders pools (subsets) clustered into different numbers of groups based on standard deviations of rmax. We cluster standard deviations of rmax into 2 and 3 different groups to show their distribution. Considering there are merely 10 unique combinations of 3 orders from a total of 5 orders, we do not incorporate clusters of 4 and 5 groups for this dataset. The results remain consistent with the theoretical prediction in Fig. 2B regardless of group numbers. Details on this dataset are in Sec. S11.

**Supplementary Figure S10:**
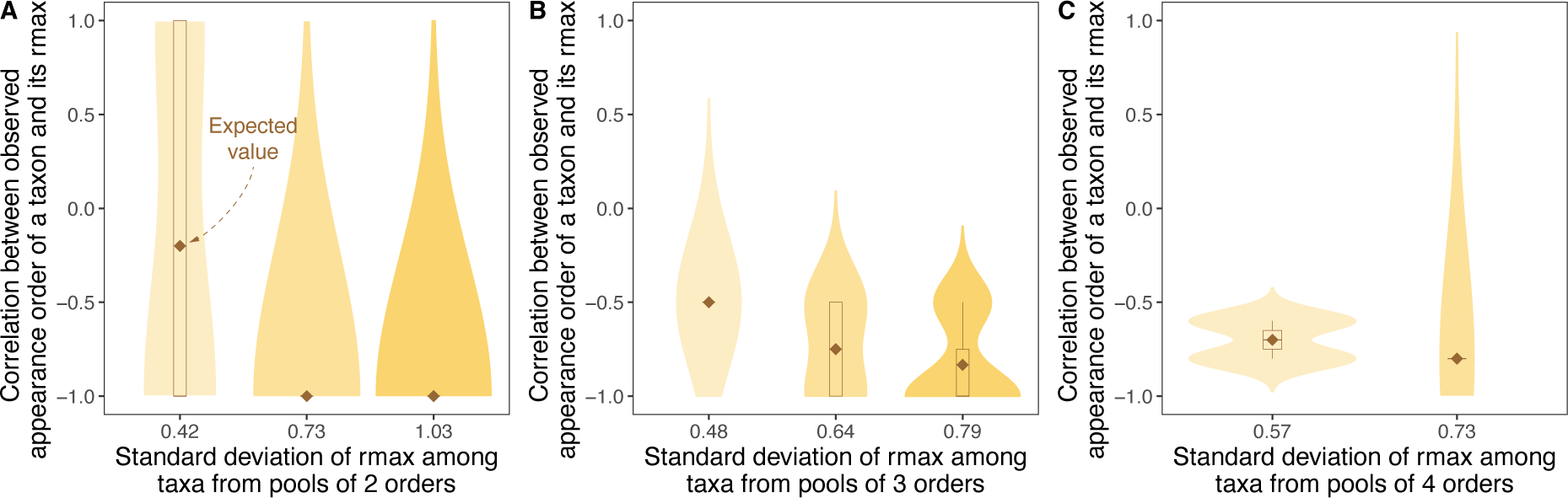
Marine bacteria pools (subsets) with different numbers of orders. We select pool sizes that are 40%, 60%, and 80% of the entire system size of 5 orders. The results remain consistent with the theoretical prediction in Fig. 2B regardless of pool sizes. Details on this dataset are in Sec. S11.

**Supplementary Figure S11:**
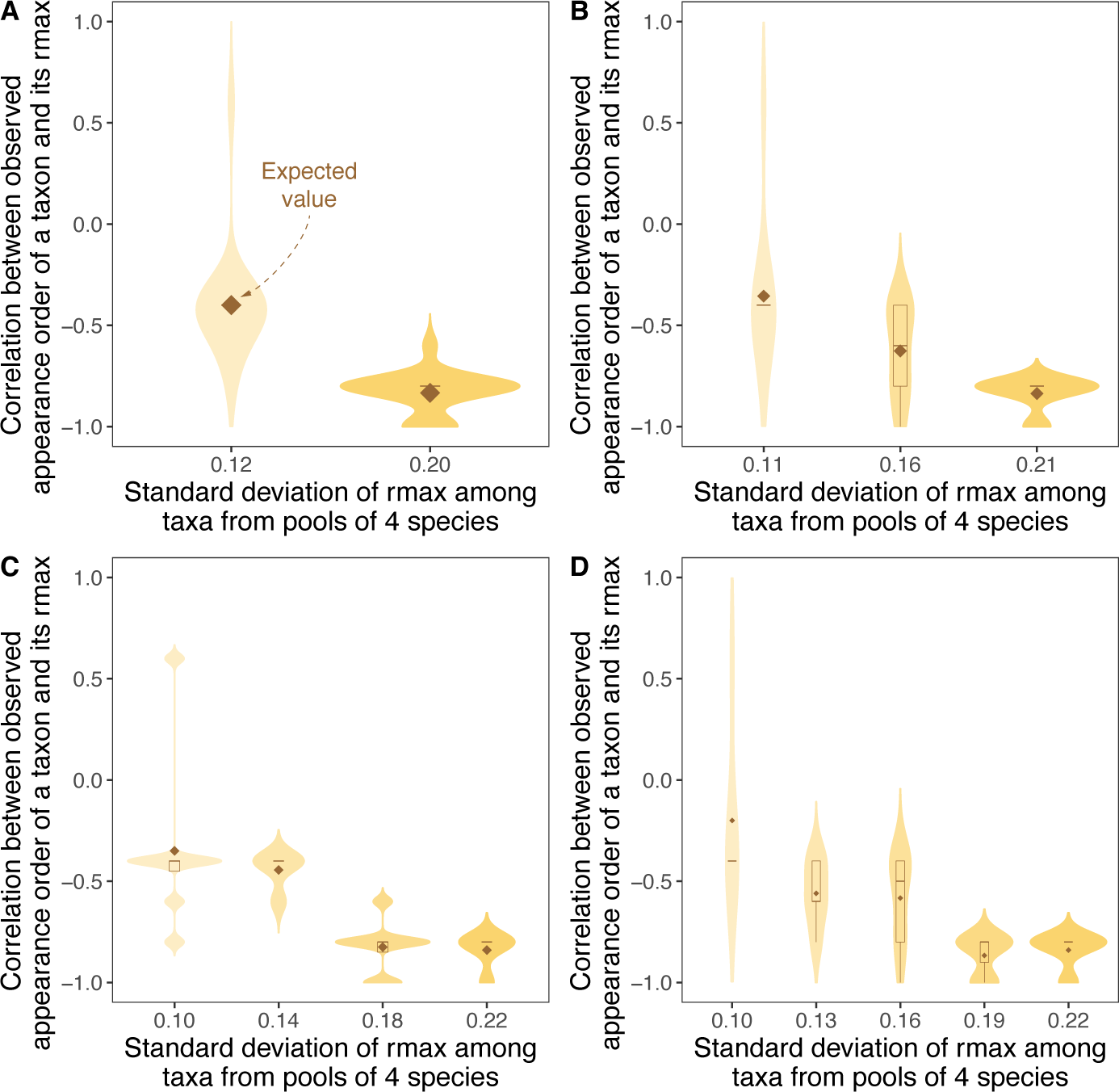
Trees 4 species pools (subsets) clustered into different numbers of groups based on standard deviations of rmax. We cluster standard deviations of rmax into 2, 3, 4, and 5 different groups to show their distribution. The results remain consistent with the theoretical prediction in Fig. 2B regardless of group numbers. Details on this dataset are in Sec. S11.

**Supplementary Figure S12:**
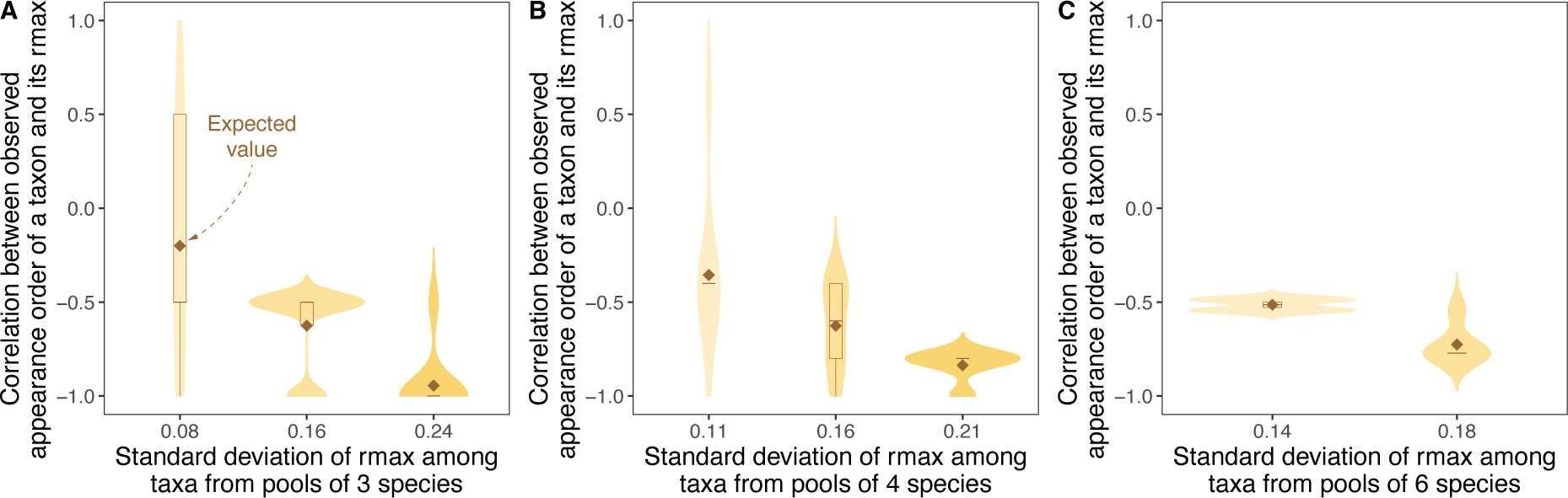
Trees pools (subsets) with different numbers of species. We select pool sizes that are 40%, 60%, and 80% of the entire system size of 7 species. The results remain consistent with the theoretical prediction in Fig. 2B regardless of pool sizes. Details on this dataset are in Sec. S11.

**Supplementary Figure S13:**
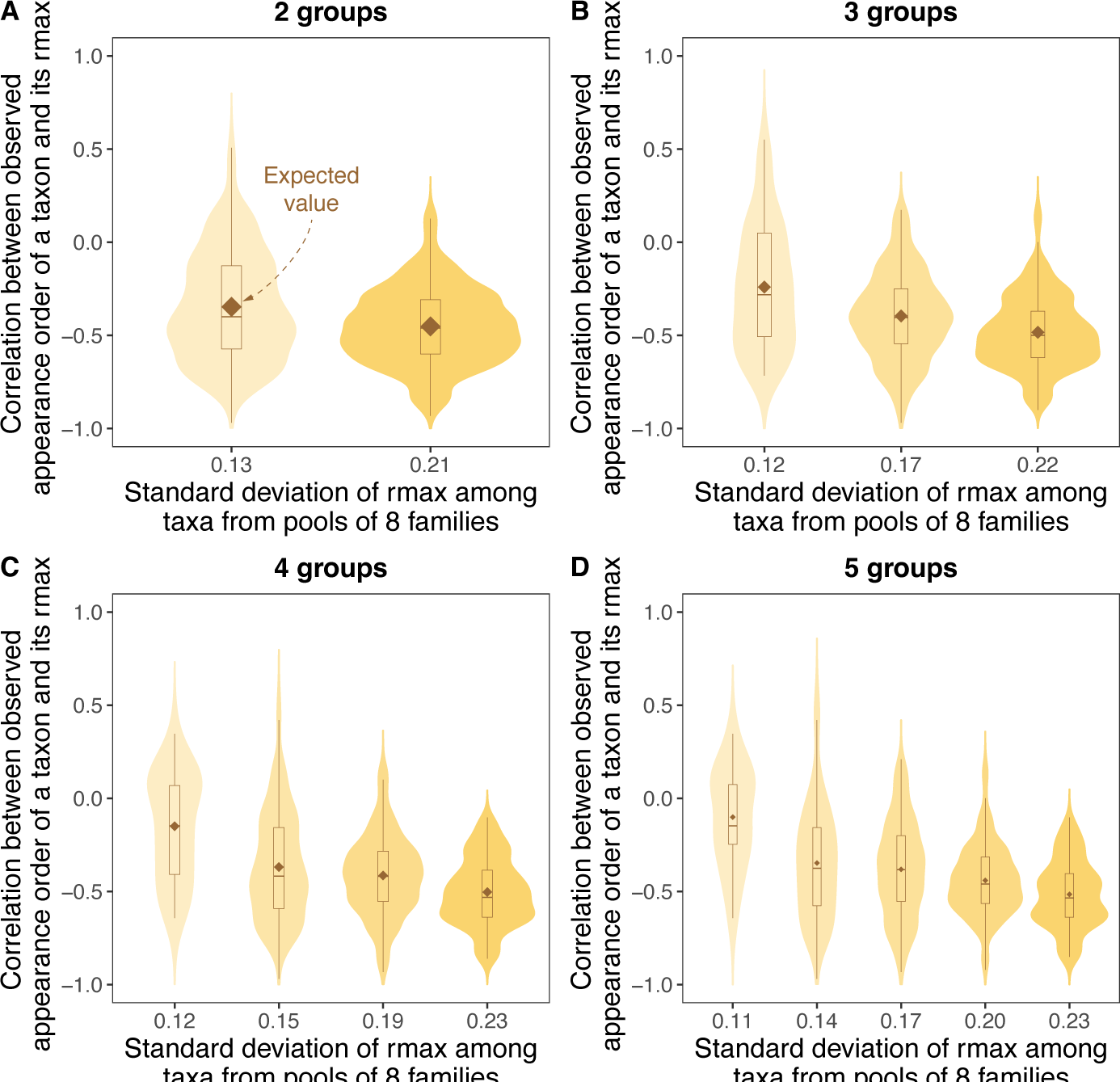
Birds 8 families pools (subsets) clustered into different numbers of groups based on standard deviations of rmax. We cluster standard deviations of rmax into 2, 3, 4, and 5 different groups to show their distribution. The results remain consistent with the theoretical prediction in Fig. 2B regardless of group numbers. Details on this dataset are in Sec. S11.

**Supplementary Figure S14:**
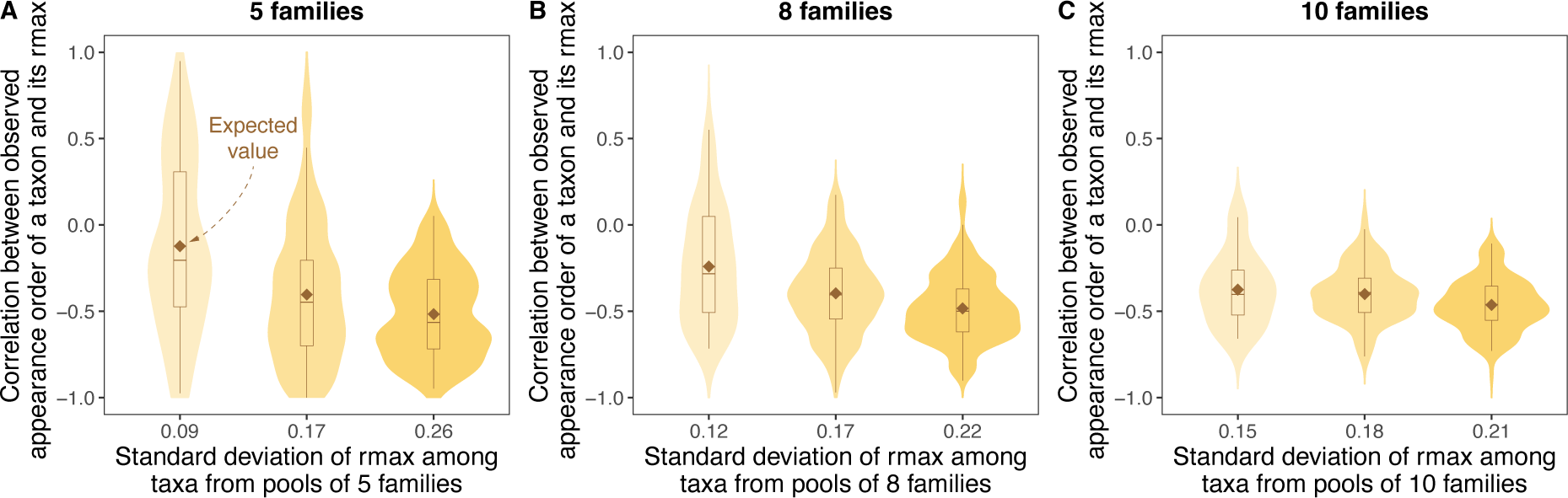
Birds pools (subsets) with different numbers of families. We select pool sizes that are 40%, 60%, and 80% of the entire system size of 13 families. The results remain consistent with the theoretical prediction in Fig. 2B regardless of pool sizes. Details on this dataset are in Sec. S11.

**Supplementary Figure S15:**
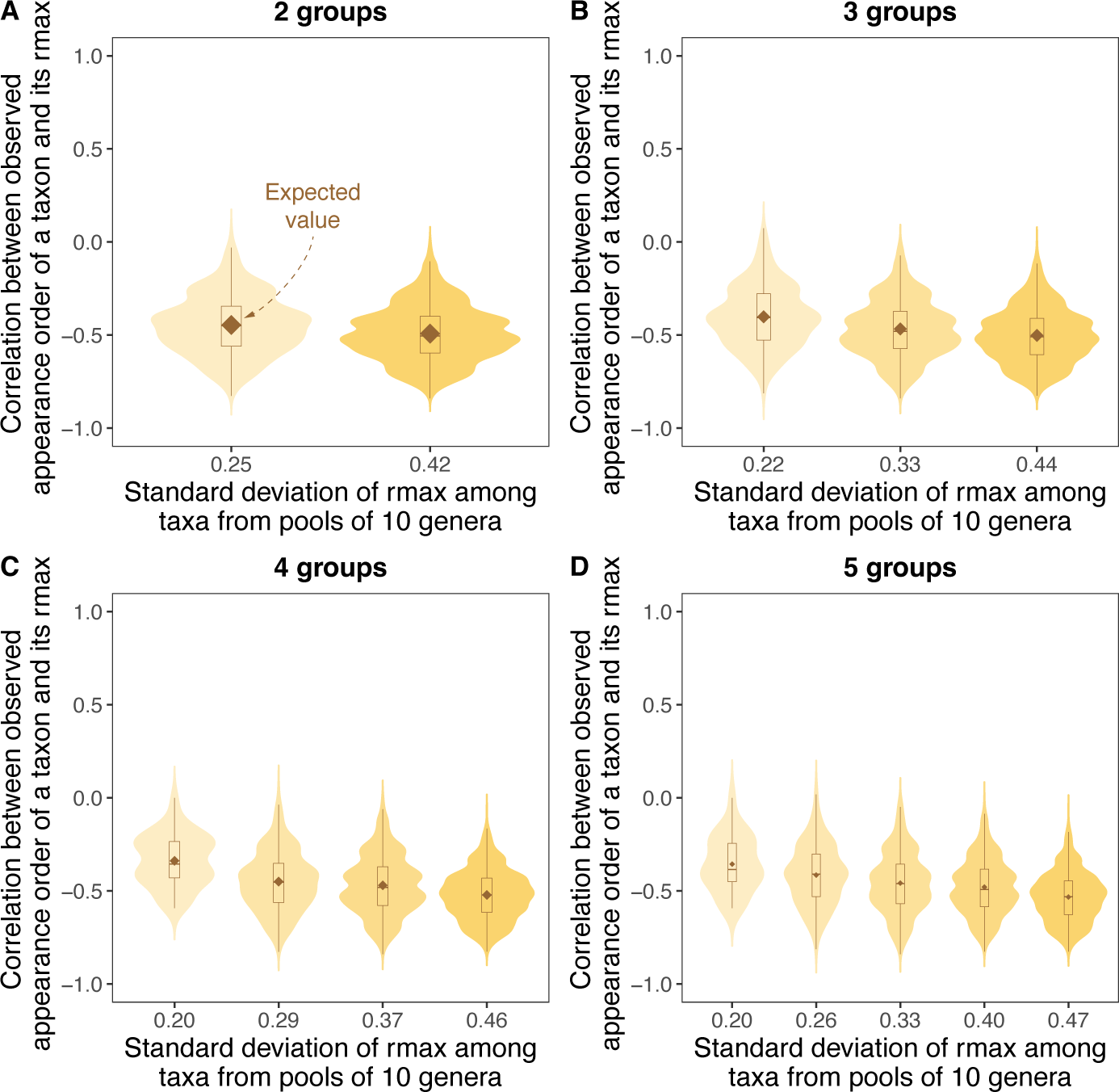
Multiyear herbivores 10 genera pools (subsets) clustered into different numbers of groups based on standard deviations of rmax. We cluster standard deviations of rmax into 2, 3, 4, and 5 different groups to show their distribution. The results remain consistent with the theoretical prediction in Fig. 2B regardless of group numbers. Details on this dataset are in Sec. S11.

**Supplementary Figure S16:**
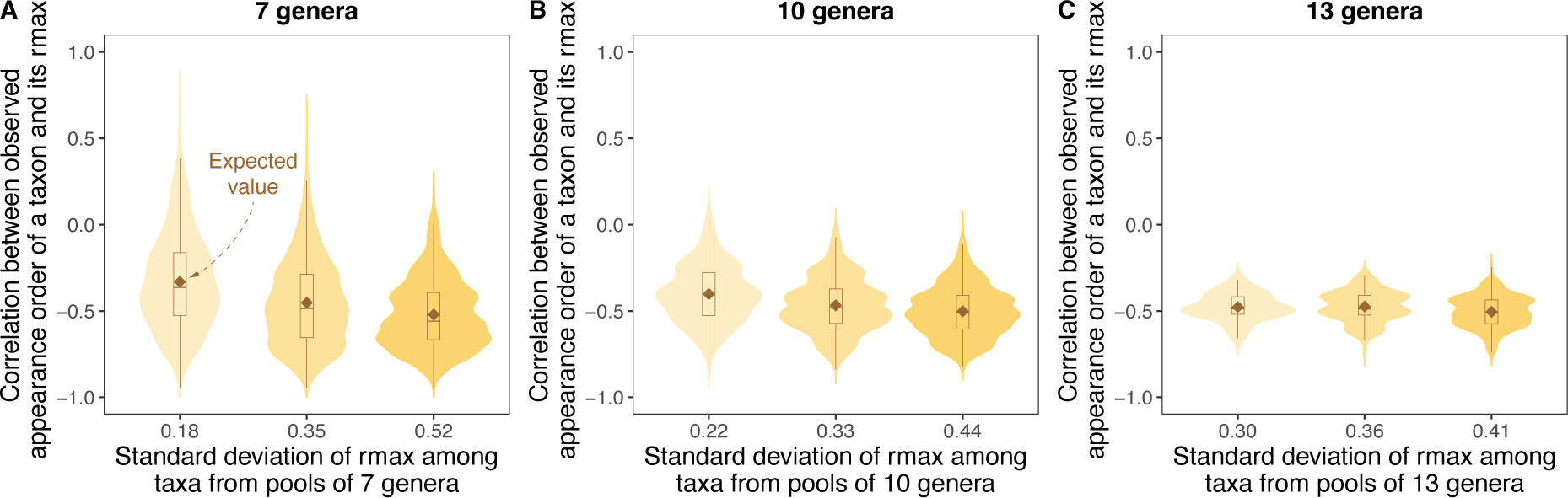
Multiyear herbivores pools (subsets) with different numbers of genera. We select pool sizes that are 40%, 60%, and 80% of the entire system size of 16 genera. The results remain consistent with the theoretical prediction in Fig. 2B regardless of pool sizes. Details on this dataset are in Sec. S11.

**Supplementary Figure S17:**
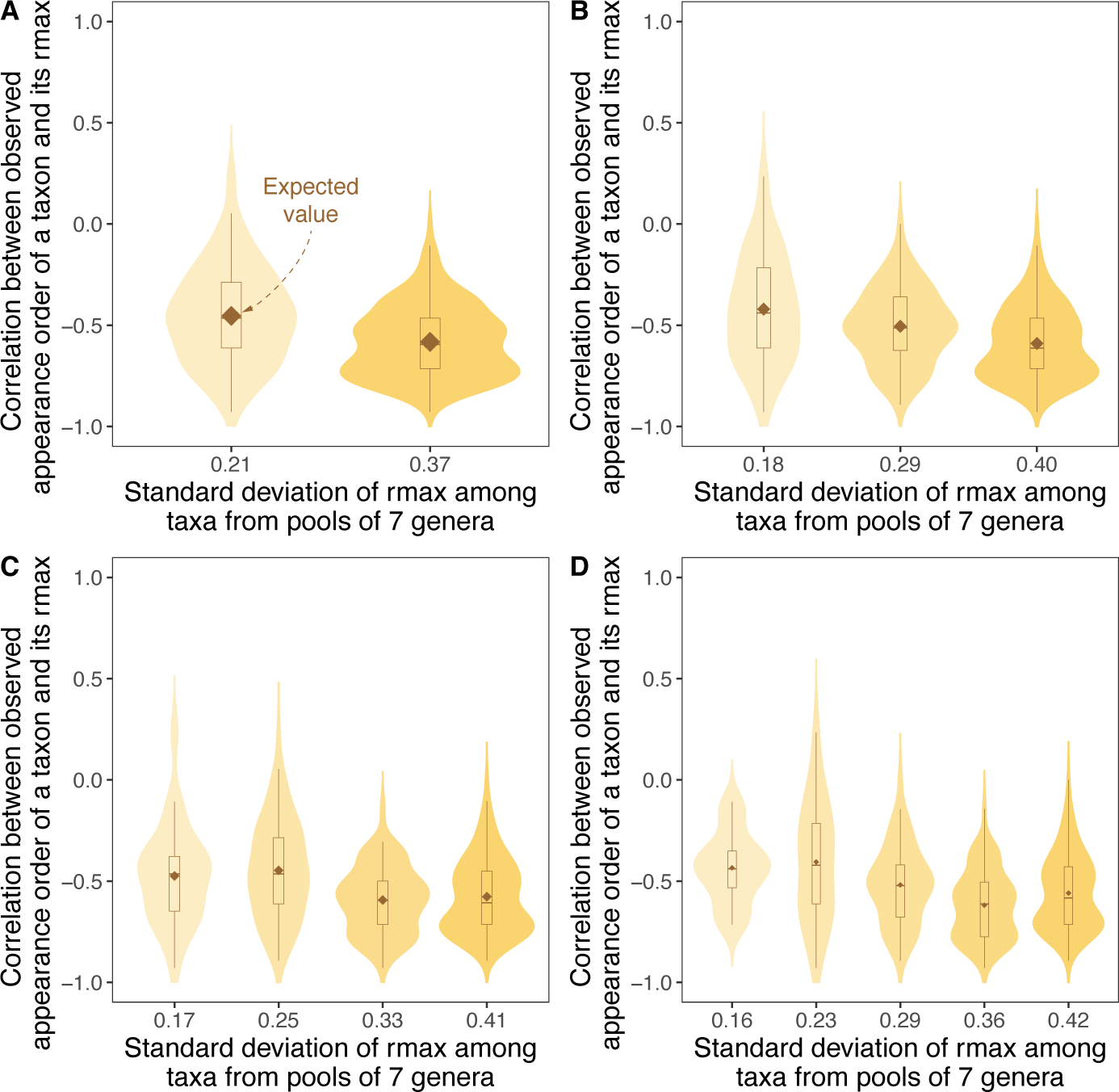
Seasonal herbivores 7 genera pools (subsets) clustered into different numbers of groups based on standard deviations of rmax. We cluster standard deviations of rmax into 2, 3, 4, and 5 different groups to show their distribution. The results remain consistent with the theoretical prediction in Fig. 2B regardless of group numbers. Details on this dataset are in Sec. S11.

**Supplementary Figure S18:**
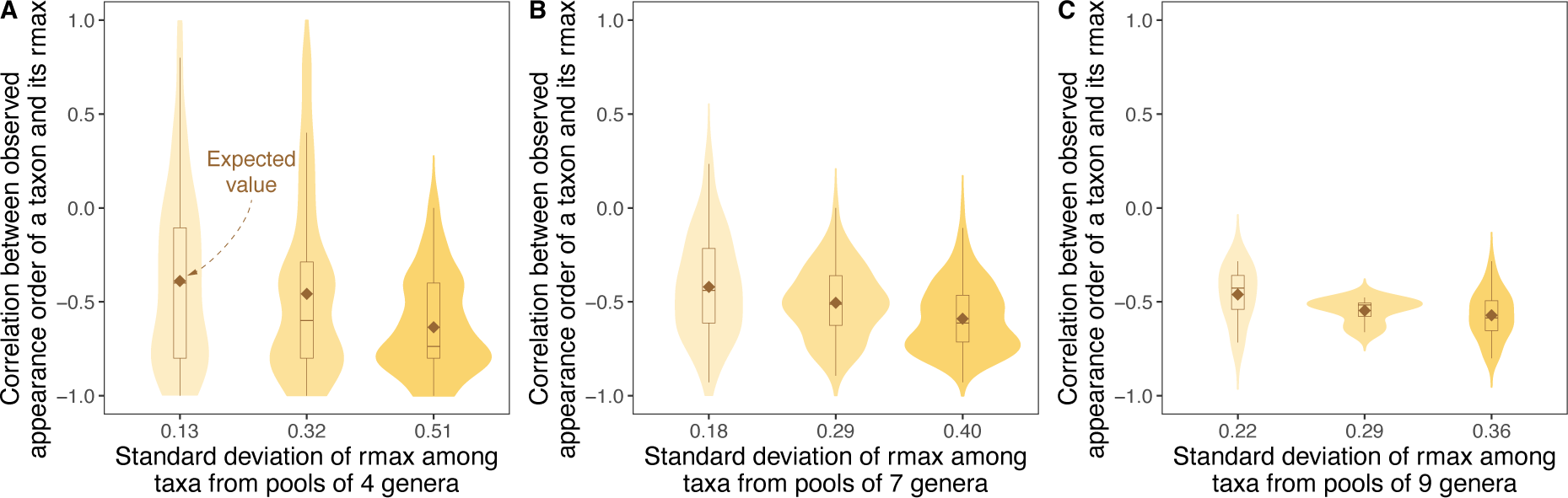
Seasonal herbivores pools (subsets) with different numbers of genera. We select pool sizes that are 40%, 60%, and 80% of the entire system size of 11 genera. The results remain consistent with the theoretical prediction in Fig. 2B regardless of pool sizes. Details on this dataset are in Sec. S11.

**Supplementary Figure S19:**
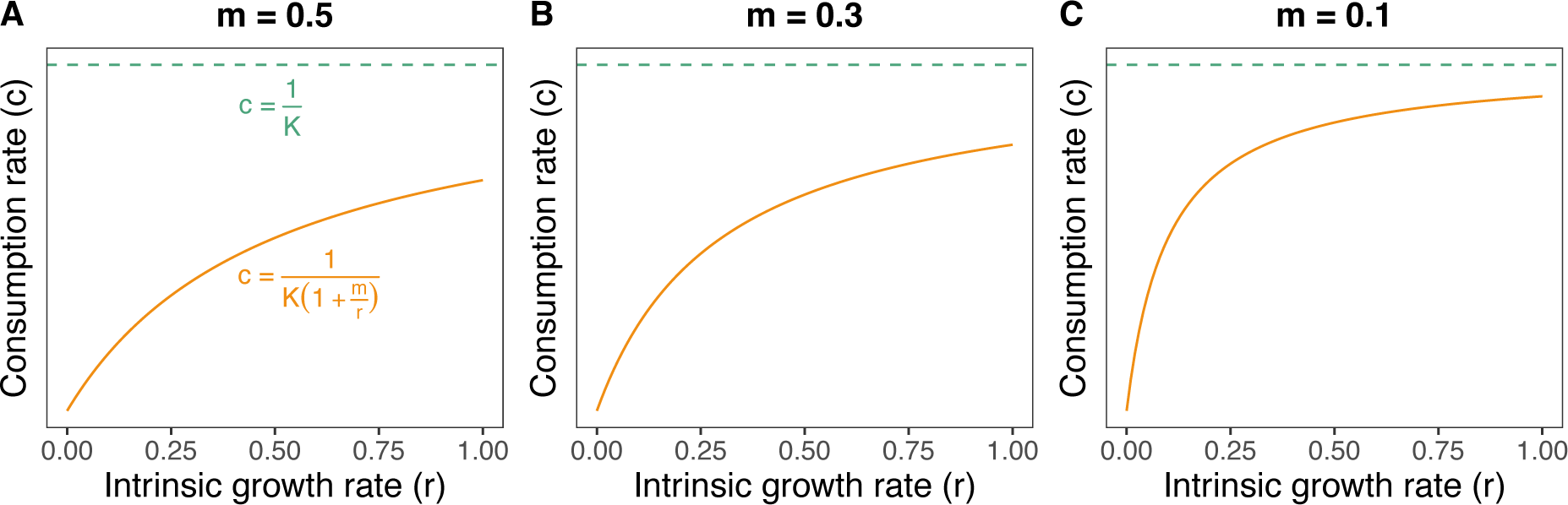
Consumption rate (c) increases monotonically with intrinsic growth rate (r) up to a maximum limit (1/K)

**Supplementary Figure S20:**
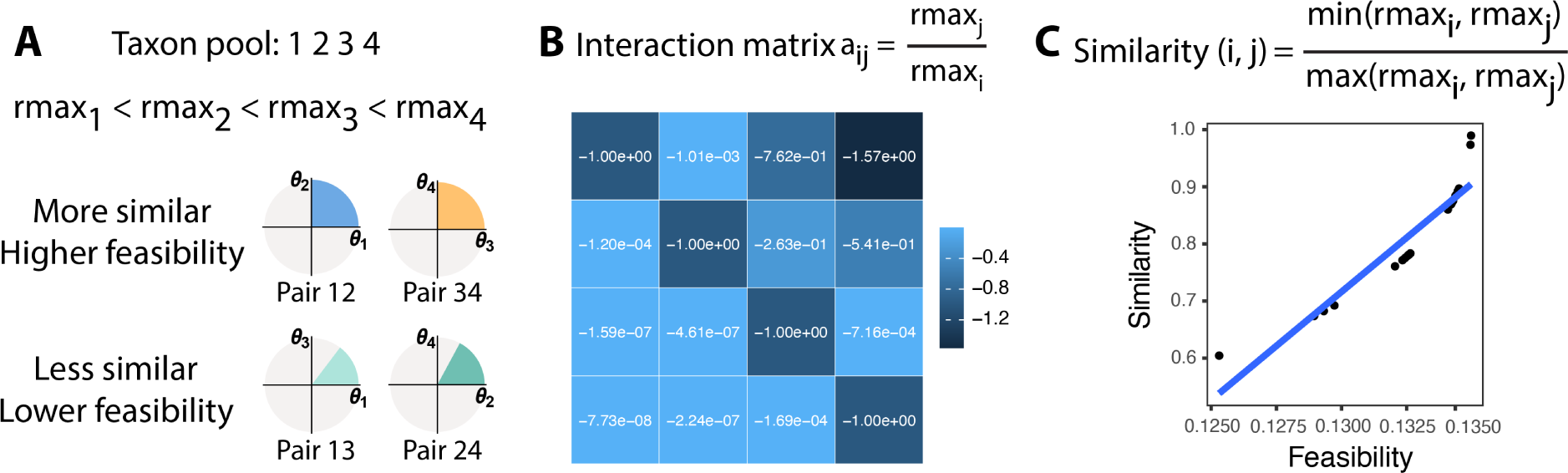
Greater similarity in growth rates among taxa enhances community feasibility. (B) For a four-taxon community, we randomly generate their body masses following a log-normal distribution with mean 0 and standard deviation 1 and then estimate their maximum growth rates (rmax) based on metabolic scaling laws. Communities formed by similar taxa tend to have higher feasibility (colored regions). The result is robust to distribution parameters. (B) The estimated interaction matrix of the four-taxon community in Panel A. (C) A confirmation of the positive relationship between similarity and feasibility. This panel shows the similarities between all pairs of a randomly generated seven-taxon community. The result is robust to community sizes.

**Supplementary Figure S21:**
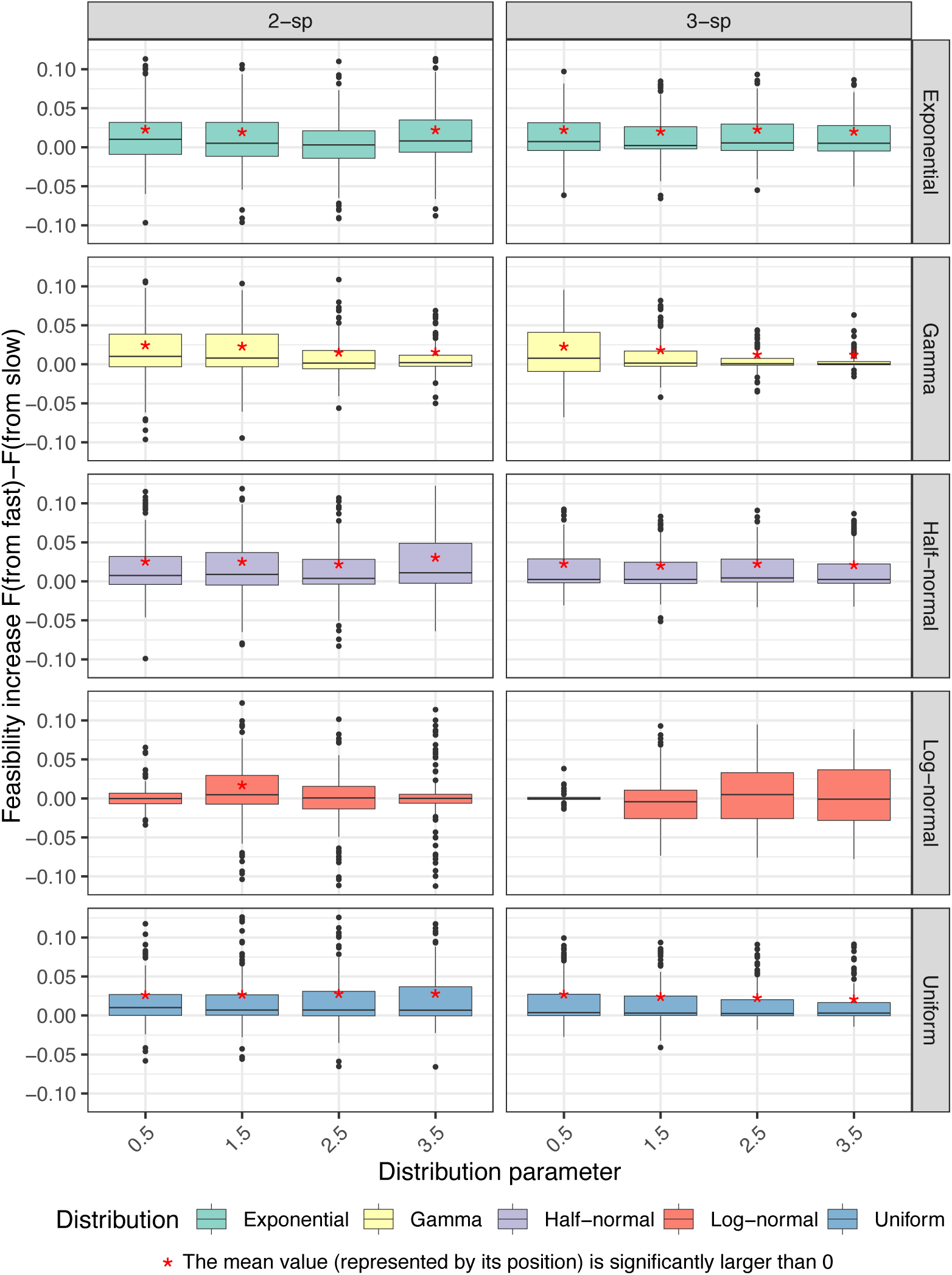
The order from fast-to slow-growing, as opposed to low- to fast-growing, is favored when the distribution deviates from log-normal distributions. When sampling body mass from exponential, gamma, half-normal, and uniform distributions, starting from fast- to slow-growing taxa is significantly more feasible than the opposite (mean*>*0, one sample t-test p-value*<*0.05, red *∗*; except for the samples in the exponential distribution with *λ* = 2.5). When sampling body mass from log-normal distributions, the randomness in small samples can lead to deviations from the perfect log-normal distribution (e.g., this holds in all cases but is particularly evident in samples with *σ* = 1.5). A minor deviation can tip the balance in the comparison between starting from fast- or slow-growing taxa, leading the system to prefer an order that starts with fast-growing taxa. This observation is based on sampling 500 communities, each with 4 taxa, for each distribution parameter.

